# Anatomy and Symbiosis of the digestive system of the vent shrimps *Rimicaris exoculata* and *Rimicaris chacei* revealed through imaging approaches

**DOI:** 10.1101/2022.04.04.486049

**Authors:** Marion Guéganton, Ouafae Rouxel, Lucile Durand, Valérie Cueff-Gauchard, Nicolas Gayet, Florence Pradillon, Marie-Anne Cambon-Bonavita

## Abstract

The shrimps *Rimicaris exoculata* and *Rimicaris chacei* are visually dominant fauna co-occurring at deep-sea hydrothermal sites of the Mid-Atlantic Ridge (MAR). Their co-existence was related to contrasted life-history traits, among which differences in their diet and reliance on chemoautotrophic symbionts at adult stage. Both shrimps are colonized by diversified chemosynthetic symbiotic microbial communities in their cephalothoracic cavity. Symbiotic association with bacteria was also evidenced in their digestive system, and the major lineages were identified through sequencing (*Mycoplasmatales* lineages mainly in the foregut and *Deferribacteres* lineages mainly in the midgut) but their clear distribution within each host species was not assessed. For the first time, we used Fluorescence *in situ* Hybridization (FISH) to visualize these lineages. Then, we described their association with digestive structures of both *Rimicaris* species. The aim was to identify possible differences between host species that could be related to their different life-history traits. For this purpose, we first developed specific FISH probes targeting *Deferribacteres* and *Mycoplasmatales* lineages identified in the digestive system of these shrimps. After signal specificity validation for each the new probe, we showed a partitioning of the bacterial lineages according to the digestive organ. Despite morphological differences between the foregut of *R. exoculata* and *R. chacei* that could be related to the adult diet, our FISH results showed overall similar distribution of digestive symbionts for the two host species. However, a more comprehensive study is needed with specimens at different life or molt stages to bring potentially host specific patterns out. Such comparative approach using FISH is now warranted thanks to our newly designed probes. These will be valuable tools to track symbiont lineages in the environment, allowing a better understanding of their relationship with their host along its life cycle, including acquisition mechanisms.

## Introduction

The Mid-Atlantic Ridge (MAR) comprises several deep-sea hydrothermal vents which shelter a dense, endemic and diversified fauna. In such environments where light does not penetrate, chemosynthesis replaces photosynthesis at the base of the trophic food web. Chemoautotrophic microbial communities ensure the primary production and the trophic resource of many species, including heterotrophic microbial communities and metazoans. They can be found as free-living communities, or associated with animal hosts (Dubilier et al., 2008; Sogin et al., 2021), and are then called holobionts (Zilber-Rosenberg and Rosenberg, 2008). This is notably the case of microbial communities symbiotically associated with the vent endemic shrimps *Rimicaris exoculata* and *Rimicaris chacei* (Williams and Rona, 1986).

The caridean shrimps *R. exoculata* and *R. chacei* co-occur at the active vents of many MAR sites (Zbinden and Cambon-Bonavita, 2020) (Fig. 1A). *R. exoculata* shrimps live in dense aggregations around active hydrothermal vents, close to the emitted fluids, and are therefore exposed to high temperatures and high concentrations of toxic compounds (Schmidt et al., 2008a, 2008b) (Fig. 1B). Juveniles of *R. exoculata* also occur within these aggregations or adjacent to them (Hernández-Ávila et al., 2021; Methou et al., 2022). *R. chacei* shrimps live at close periphery of *R. exoculata* aggregations, and are sometimes observed hiding in crevices under the rocks, or behind mussels (Methou et al., 2022). They can occasionally form small aggregations, but are found in much lower densities than *R. exoculata* adults are. In contrast, numerous juveniles of *R. chacei* live in nurseries (Methou et al., 2020; Hernández-Ávila et al., 2021) suggesting that the relatively low adult number results from a population collapse during the recruitment process of this species (Methou et al., 2022). Such drop in abundance between life stages does not seem to exist in *R. exoculata* and raises questions on the possible underlying mechanisms. The acquisition of symbiotic communities throughout each species recruitment process may play a major role in holobiont fitness (Methou et al., 2022).

**Figure 1:**
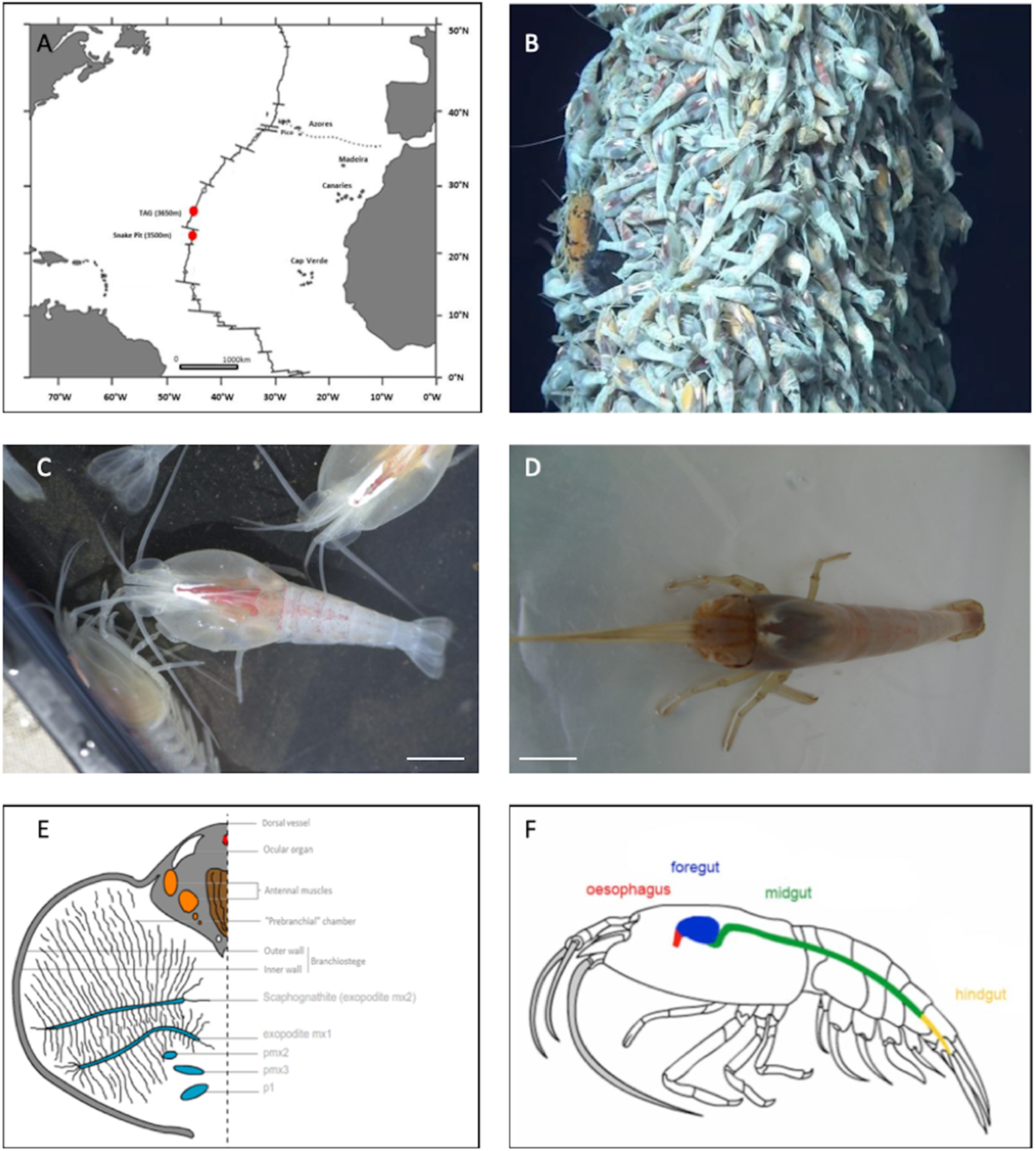
Study models. **(A)** Two main study sites on the MAR: TAG and Snake Pit. **(B)** Rimicaris exoculata aggregations on a hydrothermal vent at the TAG site (Nautile-BICOSE2, 2018). **(C)** Adult Rimicaris exoculata. **(D)** Adult Rimicaris chacei. **(E)** Transversal section of a half of the cephalothoracic cavity of R. exoculata (adapted from Segonzac et al., 1993). (**F**) Digestive system of Rimicaris sp. (adapted from Durand et al., 2009).

In adulthood, *R. exoculata* is easily distinguishable from other caridean shrimps including *R. chacei* as it has marked anatomical features (Fig. 1C, 1D). It has a reduced rostrum, its eyes have fused to form an eye plate (Williams and Rona, 1986; Komaï and Segonzac, 2008), its cephalothoracic cavity, comprising the mouthparts (scaphognathites and exopodites), is enlarged (swollen, hypertrophied) and almost closed (Van Dover et al., 1988; Segonzac et al., 1993) (Fig. 1C, 1E). *R. chacei* is somewhat different from a morphological point of view. Its cephalothoracic cavity is not as hypertrophied, since the first two pairs of chelipeds remain visible and functional (Casanova et al., 1993; Segonzac et al., 1993). Its mouthparts do not show the same morphological adaptations as those of *R. exoculata*: they are less enlarged (Apremont et al., 2018) (Fig. 1D). These morphological differences may be related to the level of development of the symbiotic bacterial communities, where more extensive communities are observed along with hypertrophied head organs (Segonzac et al., 1993).

*R. exoculata* and *R. chacei* are both colonised by three dense ectosymbiotic bacterial communities. The first described community is housed in their cephalothoracic cavity and colonises all appendages (scaphognathites, expodites and their associated setae) and the inner face of the branchiostegites (Fig. 1E). The cephalothoracic community is dominated by *Campylobacteria* (up to 90%), and *Gammaproteobacteria* (between 10-30% depending on specimens), *Alphaproteobacteria*, *Desulfobulbia* (previously named *Deltaproteobacteria*)*, Zetaproteobacteria* and *Bacteroidetes* being found in lower abundances (Zbinden et al., 2008; Petersen et al., 2010; Hügler et al., 2011; Guri et al., 2012; Jan et al., 2014; Apremont et al., 2018, Jiang et al., 2020; Cambon-Bonavita et al., 2021). These lineages form microbial mats (Zbinden et al., 2004, 2008; Corbari et al., 2008; Le Bloa et al., 2017), in which thick and long or thin and long filaments - *Campylobacteria* and *Gammaproteobacteria*, respectively - are predominant, but rod shaped bacteria (thereafter-called rods) and cocci are also observed. These microbial mats are cleared, together with the minerals and the cuticle at each molt event (every 10 days – Corbari et al., 2008). The recolonization of the cephalothorax takes place in patches on the new cuticle (Corbari et al., 2008b) and the microbial mats are then gradually formed. For *R. exoculata*, *in vivo* studies showed that these symbiotic communities play a major trophic role with direct transcuticular transfer of chemosynthetic organic matter to their host (Ponsard et al., 2013). The abundance of symbionts is lower in *R. chacei* - probably in link with a less pronounced hypertrophy of the cavity and appendages. Chelipeds of *R. chacei* are still free and, together with isotopic data, let suppose a mixotrophic regime based on symbiosis, bacteriotrophy and scavenging (Casanova et al., 1993; Apremont et al., 2018; Methou et al., 2020).

Two other microbial communities are located in the digestive system, one in the foregut and one in the midgut of both shrimp species (Fig. 1F) (Durand et al., 2009, 2015; Apremont et al., 2018). The foregut community would mainly be composed of *Mycoplasmatales* (*Firmicutes*-*Bacilli*) while *Deferribacteres* would be housed in the midgut tube. In this midgut tube, long thin “spaghetti-like” bacterial cells inserted between the microvilli of epithelial cells were observed but could not be affiliated to any bacterial lineages identified through sequencing (Durand et al., 2009, 2015; Apremont et al., 2018). Morphologically, they do not resemble described *Deferribacteres* species, which are usually small curved rods (Garrity et al., 2001). The closest relative of *Deferribacteres* lineage from the midgut of *Rimicaris* spp. is *Mucispirillum schaedleri,* isolated from rodent digestive mucus layer (Robertson et al., 2005). As for the *Deferribacteres*, the location of the *Mycoplasmatales* in the digestive system is still unclear. First, these bacteria were mostly identified in the foregut of the host using 16S rDNA approach (Durand et al., 2015) but this was not fully supported through NGS approaches (Cowart et al., 2017). *Mycoplasmatales* are heterotrophic cell wall-less bacteria usually having a reduced genome (Tully et al., 1993). *Rimicaris* spp. *Mycoplasmatales* closest relatives were identified in the hepatopancreas of the terrestrial isopod *Porcellio scaber* appearing mostly as amorphous coccoid cells (Wang et al., 2004).

Although a symbiosis has been suggested in the digestive system of *Rimicaris* shrimps, information regarding its structure and organization is relatively poorly documented, impairing a full understanding of its functioning. In farmed penaeid and caridean shrimps, the foregut is considered as one of the most complex organs to describe and understand in the animal kingdom (Ceccaldi, 1989; Štrus et al., 2019). In decapods, the digestive system is divided in three regions: the foregut, the midgut and the hindgut ((Vogt, 2021); Fig. 1F). The foregut is the anterior part of the digestive system comprising the œsophagus and the stomach. The hindgut - the terminal excretion zone - is located at the end of the abdomen (Vogt, 2021). These two regions have an ectodermic origin. Consequently, they are lined by a cuticle, which is exuviated during the molt. The third region linking the two others is the midgut, comprising the hepatopancreas and the midgut tube (Vogt, 2021). Contrary to the hindgut or the foregut, the midgut is devoid of a cuticle due to its endodermic origin, and so do not undergo molt event.

The digestive system of *R. exoculata* seems to differ from that of other crustaceans (Ceccaldi, 1989; Komai and Segonzac, 2008; Lima et al., 2016). Indeed, the œsophagus, the stomach and the hindgut are reduced, in contrast to the midgut which is long (Komaï and Segonzac, 2008; Durand et al., 2009). The stomach is reduced to a single and rather small cavity (Segonzac et al., 1993; Durand et al., 2009) (Fig. 1F). The stomach of *R. chacei*, on the other hand, is proportionally more similar to that of other caridean shrimps and more developed than that of *R. exoculata* (Segonzac et al., 1993; Komai and Segonzac, 2008, Apremont et al., 2018). This is in agreement with a mixotrophic diet involving both symbiosis and scavenging (Casanova et al., 1993; Segonzac et al., 1993; Gebruk et al., 1997, 2000; Apremont et al., 2018; Methou et al., 2020). The alimentary bolus of *R. exoculata* shrimps is mainly composed of minerals: iron sulphides and oxides, phosphate and calcium sulphate at different stages of oxidation, as well as some cuticle debris from molts (Segonzac et al., 1993). The food bolus of *R. chacei* is also full of minerals. However, it shows a somewhat different composition: in addition to minerals and cuticle debris, organic waste and organic debris can be found, confirming a mixotrophic diet (Casanova et al., 1993; Apremont et al., 2018).

In *R. exoculata*, the different microbial lineages retrieved in the digestive system seem to be resident symbionts and not part of the alimentary bolus. Indeed, even after 72 hours of fasting - emptying the digestive system - they are still present (Durand et al., 2009). The foregut, covered by a cuticle, molts every 10 days (as the cephalothorax does - Corbari et al., 2008), contrary to the midgut. Inter molt and inter generation transmissions of lineages affiliated to *Mycoplasmatales* and *Deferribacteres* remain enigmatic, as are their potential roles. No *Deferribacteres* and only one *Mycoplasmatales* affiliated OTU (Hügler et al., 2010; Flores et al., 2011) was identified from the shrimp’s aggregations environment. The level of similarity between *Deferribacteres* 16S rDNA gene sequences, whatever the site of sampling, is higher than 99%, suggesting a single phylotype, which may suppose a vertical transmission (Durand et al., 2015). However, this lineage was not found in microbial communities associated with broods (Guri et al., 2012; Cowart et al., 2017; Methou et al., 2019), which would rather indicate horizontal transmission. Until now, the key questions related with the acquisition of symbionts at the juvenile stage and after each molt, the overall functioning of the digestive system and symbiont niche distribution are still debated.

To better describe symbiont niche distribution, to infer their potential roles and the overall functioning of holobionts, we used fluorescent *in situ* hybridization (FISH) approaches with newly designed molecular probes targeting specifically rRNA sequences of these digestive symbiotic lineages. This approach allows both native symbiotic cell visualization (morphology and localization on host organ/cells) and phylogenetic identification (Amann et al., 1990, 2001). Yet, *in situ* observations of *Deferribacteres* and *Mycoplasmatales* lineages in *Rimicaris* spp. tissues have failed so far, as no specific molecular probe but the universal Eub338 bacterial probe, ever gave any positive signal (Durand et al., 2009, 2015).

In this study, we designed and validated new FISH probes targeting specifically the *Deferribacteres* and the *Mycoplasmatales* lineages found in the digestive system of both *Rimicaris* spp. These probes allowed us to visualize and identify digestive symbiont lineages in adult shrimps. The combination of FISH approaches with SEM imaging of the digestive structures of both shrimp species was used to answer the following questions: 1) What is the distribution and abundance of digestive symbiotic lineages in each *Rimicaris* species? 2) Are symbionts associated with specific morphological structures in each host, possibly reflecting niche partitioning? 3) Can symbiont distribution and abundance in their respective hosts be related to specific ecological traits of the holobionts (e.g. nutrition type)?

## Materials and Methods

### Sampling and specimens conditioning on board

Samples were collected at two vent fields along the MAR: TAG (26°8 “N-44°50 “W, 3650 m depth) and Snake Pit (23°22 ’’N; 44°57’’W, 3460 m depth) during the BICOSE2 cruise (26 January to 10 March 2018, DOI http://dx.doi.org/10.17600/18000004). The specimens were caught in shrimp aggregations using the suction sampler of the HOV (Human Operated Vehicle) Nautile operated from the R/V *Pourquoi pas*?

Once on board, shrimps were dissected under sterile conditions to recover different anatomical parts: the branchiostegites (LB) and the scaphognathites (Sc) in the cephalothoracic cavity, and the digestive system comprising the foregut and midgut. The different organs were immediately fixed for the FISH study in a 3% formalin seawater solution for 3 hours. This step is essential to ensure proper fixation for cell integrity. Samples were then rinsed three times with a phosphate buffered saline solution (PBS) and stored in a PBS/Alcohol (1:1) solution at -20°C (Durand et al, 2009). Some digestive tissues were fixed in a 2.5% glutaraldehyde solution (16 hours at 4°C), and then rinsed and stored at 4°C in a solution containing a biocide to avoid bacterial development [filtrated sea-water with 0,44 g/L of NaN_3_ at pH 7,4] until use for scanning electron microscopy (SEM) observations. In addition, whole specimens were frozen at -80°C for later dissections at the laboratory (Table 1).

**Table 1:** Samples selected for the different approaches: SEM, FISH and dissection under binocular microscope.

| Vent fields | Foregut | Midgut tube | Scaphognathites |
| --- | --- | --- | --- |
| TAG | 8 *R. exoculata* → FISH<br>1 *R. chacei* → FISH<br>2 *R. exoculata* and 1 *R. chacei* → anatomy | 3 *R. exoculata* → FISH<br>1 *R. chacei* → FISH | 1 *R. exoculata* → FISH |
| Snake Pit | 10 *R. exoculata* → FISH<br>1 *R. chacei* → FISH<br>2 *R. exoculata* → SEM | 2 *R. exoculata* → FISH<br>1 *R. chacei* → FISH |  |

### In silico design and validation of new FISH specific probes

The FISH method is based on the use of specific molecular probes linked to a fluorescent dye, which are complementary to a region of the 16S or 23S rRNA target molecule, directly inside the native ribosomes of fixed microbial cells. The new probes targeting specifically *Mycoplasmatales Rimicaris* symbiont lineages (Durand et al., 2009, 2015) and *Deferribacteres Rimicaris* symbiont single lineage (Durand et al., 2009, 2015) were first designed *in silico*, based on *Rimicaris* symbionts 16S rDNA sequences available in the literature (Durand et al., 2009, 2015; Apremont et al., 2018). These sequences were aligned with the MUSCLE (MUltiple Sequence Comparison by Log-Expectation) algorithm (Edgar, 2004) in Geneious software v9 (Kearse et al., 2012). This software allows to highlight homology and evolutionary relationships between the studied sequences and thus, to select rRNA molecule regions specific of the digestive symbiont lineages. Literature data on molecular probes design (Behrens et al., 2003) were used to select 16S rRNA molecule regions known to be accessible for probe hybridization, *i.e*. avoiding complex 3D *in situ* conformations of the rRNA molecule or regions showing interactions with ribosomal proteins. Then, the physical properties of the designed probes were estimated to avoid hairpin structure formation, cross-hybridization, self-dimer formation, and to check their melting temperatures (TM), using the Geneious Primer Design tool and the Oligo Calc software (Kibbe, 2007). Finally, the complementarity of these designed probes with rRNA gene sequences of non-targeted microorganisms was evaluated more largely using BLAST (Altschul et al., 1990) with the NCBI and Silva138 databases (test probes v3.0).

### Sample preparation for Fluorescence In Situ Hybridization procedures

Tissue sections were prepared by embedding dissected organs (foregut, midgut tube and scaphognathites) in polyethylene glycol distearate-1-hexadecanol (9 : 1) resin (Sigma, St., Louis, MO) after progressive dehydration and soaking into the resin (PBS – ethanol series at ambient temperature and ethanol – resin series at 40°C) (Duperron et al., 2007). After solidification of the resin, blocks containing organs were stored at -20°C until trimming. According to the sample, 8-10 µm transversal tissue sections were obtained with a RM 2255 microtome (Leica Biosystems, Nussloch, Germany) and placed on coated slides (Menzel-Gläser Superfrost® Plus, USA). Before hybridization, resin was removed with ethanol (3x5min in 96° ethanol) and tissues were rehydrated (5 min in 70° ethanol).

### New FISH probes optimal hybridization condition determination

After *in silico* validation, optimal probe hybridization conditions were determined using host tissues known to harbour (e.g. digestive) or not (e.g. scaphognathites) these target symbiont lineages (Table 1). GAM42a targeting *Gammaproteobacteria* (Manz et al., 1992) and Epsy549 targeting *Campylobacteria* (Lin et al., 2006) were used as positive controls on scaphognathites when testing our new probes to check the correct hybridization procedure effectiveness (Table 2). The universal probe Eub338-I targeting most Eubacteria (Amann et al., 1990) was used as a general positive (co-)hybridization control (Table 2). All probes were synthesized by Eurofins Genomics (Ebersberg, Germany) and were labelled with either Cyanine 3 or Cyanine 5 dyes (Table 2).

**Table 2:**
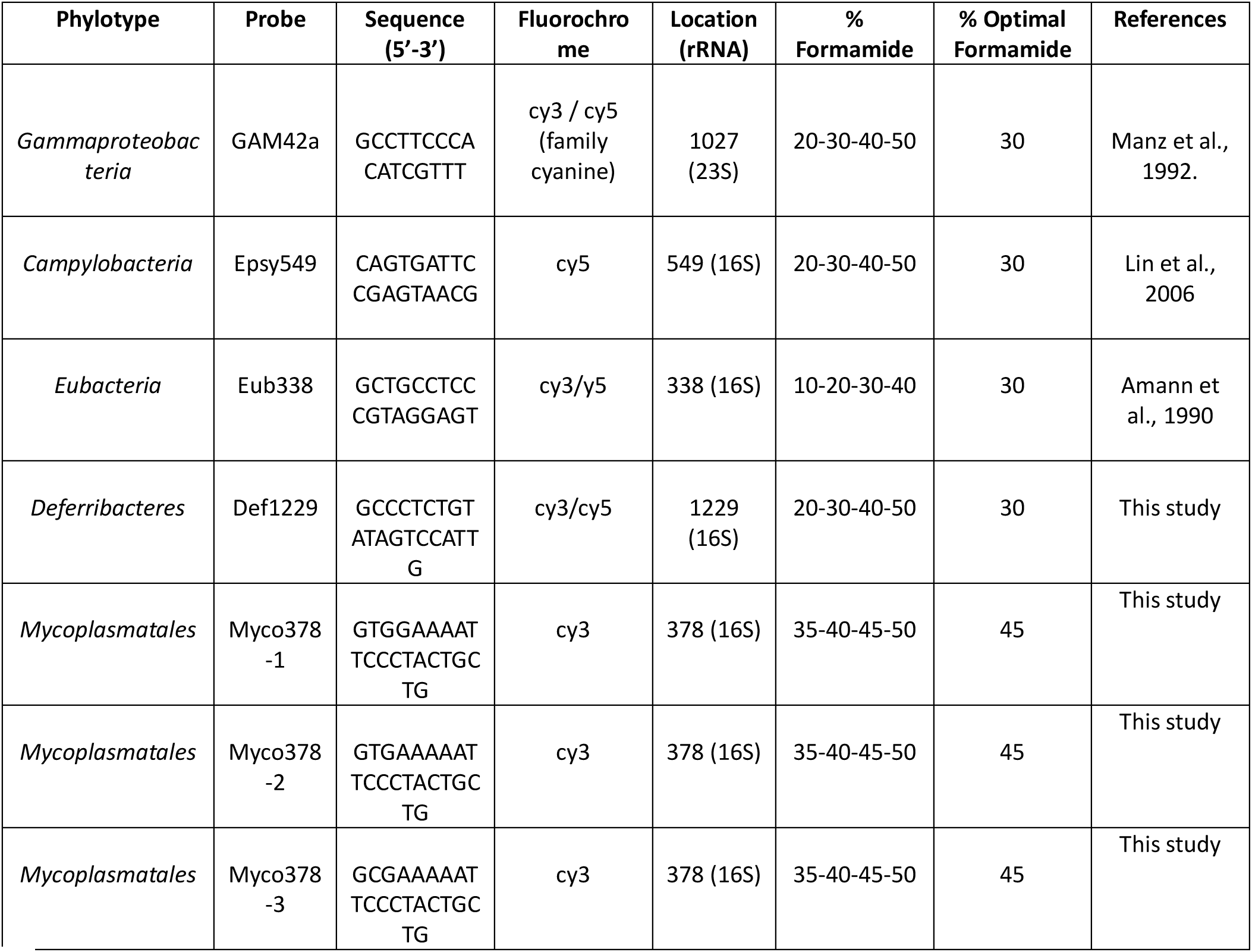
Probes used for this study.

Appropriate stringency conditions were determined using various combinations of formamide concentrations in the hybridization buffer and hybridization temperatures, both denaturants that aim to relax, but not alter the ribosome structure, enabling effective probe hybridization. Sections were soaked in reaction mix containing probes (each at 0,5µM final concentration) in a hybridization buffer [0,9M NaCl, 0,02M Tris-HCl [pH 7.5], 0,01% [w/v] sodium dodecyl sulphate (SDS), 20%, 30%, 35%, 40%, 45% or 50% deionized formamide (see Tab 3.)], and incubated for 3 hours at 46°C or 48°C (see Table 3)]. Sections were washed for 15 min or 30 min, at a slightly higher temperature (48°C or 50°C) than the hybridization temperature applied (Table 3), in a washing buffer adapted to the formamide concentrations [0,215M, 0,102M, 0,07M, 0,046M, 0,03M or 0,018M NaCl respectively for 20%, 30%, 35%, 40%, 45% or 50% formamide, 0,02M Tris-HCl [pH 7.5], 0.005M EDTA [pH 8] and 0,01% [w/v] SDS]. Then, they were briefly rinsed twice with distilled water, once at washing temperature then at room temperature. Finally, sections were mounted on slides with SlowFade™ Gold antifade reagent with DAPI (Invitrogen). Observations on hybridized tissues were made using a Zeiss Imager.Z2 microscope equipped with the Apotome.2 sliding module and Colibri.7 light technology (Zeiss, Oberkochen, Germany). The micrographs were analyzed using the Zen software (Zeiss). DIC (Differential interference contrast Differential interference contrast) was used to better visualize the host tissues.

**Table 3:** Conditions used in stringency tests performed for probes validation.

| Conditions/<br>Samples | Adult<br>*scaphognathites* | Adult<br>*Midgut tubes* | Adult<br>*scaphognathites* | Adult<br>*foreguts* |
| --- | --- | --- | --- | --- |
| Formamide in<br>hybridization<br>buffer (%) | 20% - 30% - 40% - 50% | 20% - 30% - 40% - 50% | 35% - 40% - 45% - 50% | 35% - 45% |
| Hybridization<br>temperature (°C) | 46°C | 46°C | 46°C and 48°C | 46°C |
| Washing<br>temperature (°C) | 48°C | 48°C | 48°C and 50°C | 48°C |
| Washing time<br>(min) | 15 min | 15 min | 15 min and 30 min | 30 min |
| Probes | Def1229 | Def1229 | Myco378-1<br>Myco378-2<br>Myco378-3 | Myco378-1<br>Myco378-2<br>Myco378-3 |

### Scanning Electronic Microscopy (SEM)

Samples fixed in glutaraldehyde were first dehydrated in ethanol series (10% to 100% in 8 steps). Dehydrated samples were placed in a perforated box, critical-point dried (Leica EM CPD300), sticked on carbon glue that was placed on a plot and then coated by gold-sputtering (60% gold/ 40% Palladium, Quorum Technologies SC7640). SEM observations were performed using a Quanta 200 MK microscope (FEI, Hillsboro, OR) and images were taken with the SCANDIUM acquisition program (Soft Imaging System, Munster, Germany).

## Results

### Probe sequences determination

As 16S rDNA gene sequences revealed four distinct phylotypes (Durand et al., 2010, 2015; Apremont et al., 2018), three probes were designed to cover the diversity of *Mycoplasmatales* lineages identified in *R. chacei* and *R. exoculata* foregut, namely Myco378-1, -2 and -3. A single probe was designed for the unique *Deferribacteres* phylotype, namely Def1229 (Table 2). The probes designed were: Myco378-1 (5’GTGGAAAATTCCCTACTGCTG’3), Myco378-2 (5’GTGAAAAATTCCCTACTGCTG’3), Myco378-3 (5’ GCGAAAAATTCCCTACTGCTG’3) and Def1229 (5’GCCCTCTGTATAGTCCATTG’3) (Table 2). The location of the three probes Myco378 (nucleotide position 378-398 according to *Mycoplasmatales* 16S rDNA sequence) on the 16S rRNA sequence is very close to the location of the general Eubacteria probe (nucleotide position 338-358 according to *E. coli* 16S rDNA sequence) but do not overlap and should then allow a co-hybridization.

### Determination of optimal stringency conditions for the hybridization of the probes Def1229, Myco378-1, Myco378-2 and Myco378-3

In order to ensure the specificity of the probes, tests were performed on transversal sections of different organs from several adult specimens, as none of the associated shrimp lineages could ever be grown in laboratory (Zbinden and Cambon-Bonavita, 2020). The midgut tube is supposed to be colonized by *Deferribacteres* according to previous sequencing results, and the foregut is supposed to host *Mycoplasmatales.* Scaphognathites of the cephalothoracic cavity were used as a control for non-specific hybridization since *Deferribacteres* or *Mycoplasmatales* were never detected using DNA sequencing approaches.

#### Specificity and stringency conditions of Def1229 probe hybridization

The Def1229 probe was tested on multiple midgut tube sections of an adult *R. exoculata*. Regardless of the section of the midgut tube, hybridization with the Def1229 probe always highlighted thin filamentous cells inserted between the microvilli of the midgut tube cells (Fig. 2A-E). Moreover, varying formamide concentration in the hybridization buffer had no impact on the hybridization signal. In fact, whatever the stringency conditions (20% to 50% formamide), a high fluorescence signal intensity was detected with Def1229 in the midgut tube (Fig. 2A-E, Supplementary Table 1). On sections of adult *R. exoculata* scaphognathites used as negative control, whatever the stringency (20% to 50% formamide), no fluorescence was ever detected with the Def1229 probe (Supplementary Fig. 1, Supplementary Fig. 2, Supplementary Fig. 3, Supplementary Table 1).

**Figure 2:**
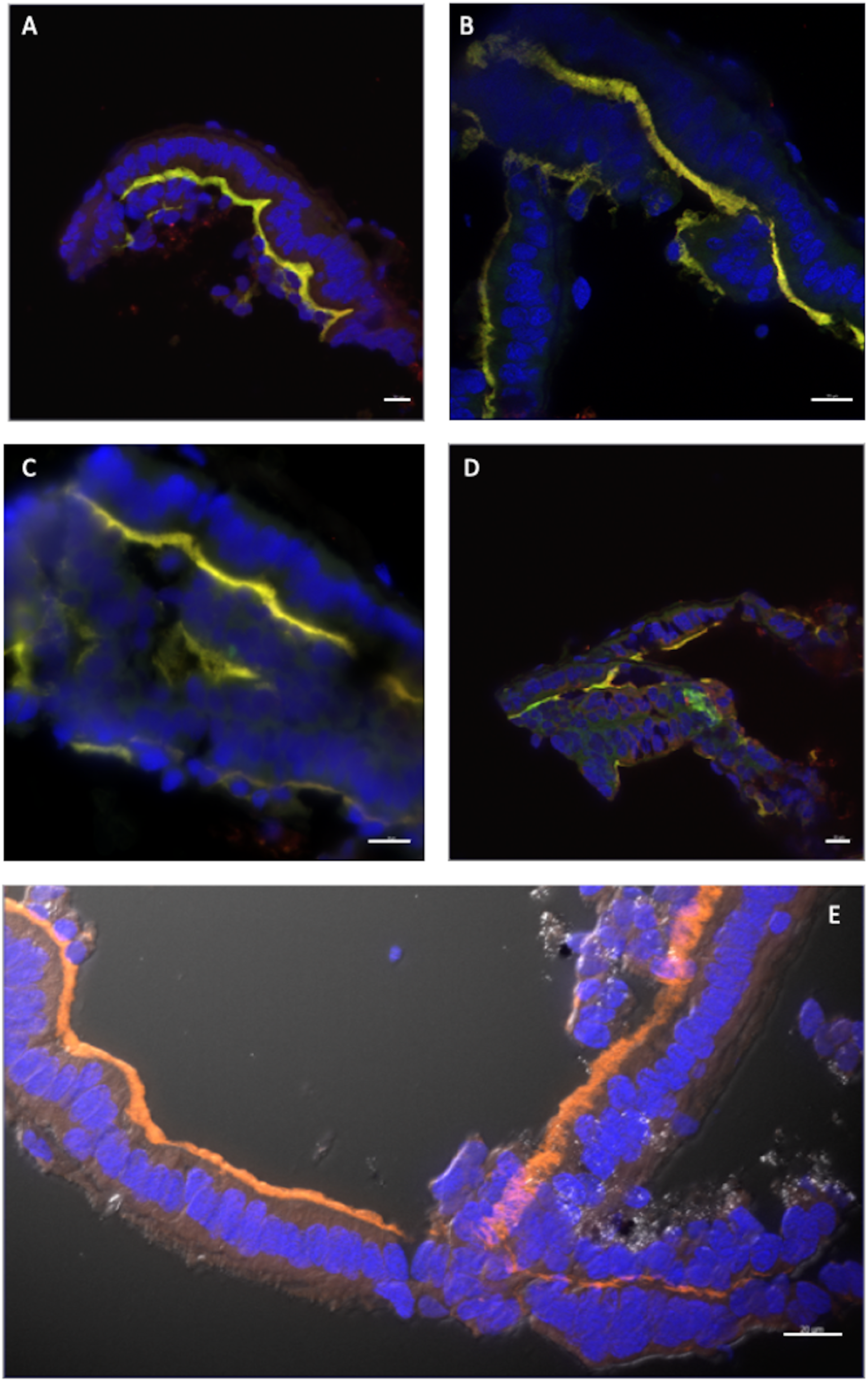
Stringency tests with Def1229. **(A-E)** represent transversal sections through the midgut tube of a R. exoculata adult hybridized with Eub338-cy3 (green) /Def1229-cy5 (red) **(A-D)** or with Def1229-cy3 (orange) only **(E)**. Double Hybridizations appear in yellow-green. Tissue cell nuclei are labeled with DAPI (blue). Formamide concentration in hybridization buffer was 20% **(A)**, 30% **(B, E)**, 40% **(C)**, 50 % **(D)**. Pictures in **(A, B, D, E)**, were taken with Apotome and with Z-stack, and with DIC (Differential interference contrast) for **(E)** only. Scale bars = 20 µm.

In order to further assess Def1229 specificity, co-hybridizations with probes Eub338, Epsy549 and GAM42a were carried out on adult midgut tubes and scaphognathites (Supplementary Fig. 1, Supplementary Fig. 2, Supplementary Fig. 3, Supplementary Table 1). These co-hybridizations confirmed the previous results. Indeed, on the scaphognathite sections, large filamentous bacteria (*Campylobacteria*), thin filamentous bacteria (*Gammaproteobacteria*), coccoids and rods were only detected respectively with the probes Epsy549, GAM42a and Eub338 whatever the stringency conditions (20% to 50% formamide) (Supplementary Fig. 1, Supplementary Fig. 2, Supplementary Fig. 3, Supplementary Table 1). On the contrary, bacteria attached to the midgut tube wall cells were only revealed by Eub338 and Def1229 probes (Fig. 2A-D, Supplementary Table 1). These observations allowed us to validate the Def1229 probe and its specificity whatever the stringency condition used.

By convention, when formamide concentration has no effect on probe stringency, hybridization buffers containing 30% formamide are retained (in agreement with the optimal protocols described for universal probes: hybridization at 46°C for 3 hours and washing at 48°C for 15 minutes). The Def1229 probe is therefore specific to *Deferribacteres* of the shrimp *R. exoculata*.

#### Specificity and stringency conditions of Myco378-1, Myco378-2 and Myco378-3 probes hybridization

To confirm the specificity of the 3 probes, first tests were performed on *R. exoculata* adult foregut sections, all along the organ from the œsophagus toward its posterior part, using hybridization buffer with 30%, 35%, 40% and 45% formamide and a hybridization temperature of 46°C (Fig. 3A-C). On a part of the foregut, many small rods were identified with each Myco378 probe (with or without co-hybridization with Eub338 probe) using 30% formamide in the hybridization buffer. A higher percentage of formamide showed a better signal on the rods (intense signal with 45% formamide) (Fig. 3C). In co-hybridization with Eub338 probe (45% formamide), these foregut rods were revealed by strong and intense fluorescence with both probes in all experiments. These rods were never hybridized with any of the probe Epsy549, GAM42.

**Figure 3:**
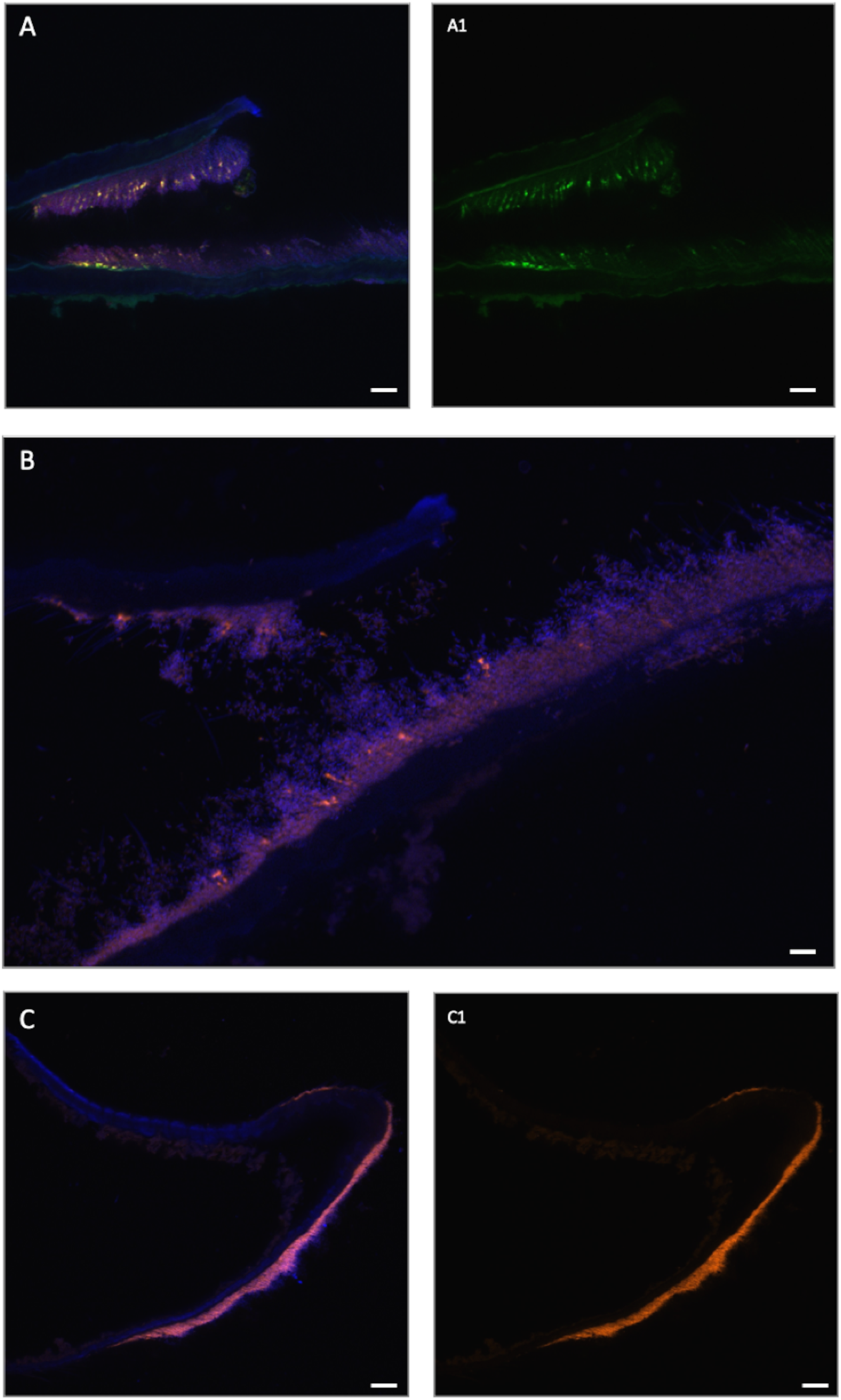
Stringency tests with Myco378-1. **(A-C)** represent sections through the œsophagus of R. exoculata adult hybridized with Myco378-1-cy3 only (yellow-green) in **(A)**, orange-pink in **(B, C)**). Tissue cell nuclei are labeled with DAPI (blue). Formamide concentration in hybridization buffer was 35% **(A, B)**, 45% **(C)**. Pictures in were taken with Apotome. Pictures **(A1, C1)** represent the same photographs as **(A, C)** without DAPI. Scale bars = 20 µm.

*R. exoculata* adult scaphognathite sections were used to perform the specificity tests with the Myco378 probes. The tests were first performed with Epsy549, GAM42a, or Eub338 co-hybridized with Myco378-1, Myco378-2 or Myco378-3 probes (30-35-40-45-50% formamide in buffer) (Supplementary Fig. 4, Supplementary Fig. 5, Supplementary Table 2). None of the Myco378 probes labelled *Gammaproteobacteria* while co-hybridized GAM42a probe properly did. Each Myco378 probe hybridized slightly *Campylobacteria* cells (largest filamentous bacteria, co-hybridized with Epsy549 probe). This faint hybridization signal decreased when a higher percentage of formamide (i.e. higher stringency) was applied to reduce non-specific hybridization. These non-specific hybridizations were lower at 40% to 50% formamide for Myco378-1, at 45% to 50% formamide for Myco378-2 probes, and at 50% formamide for Myco378-3 probe. Other tests with higher hybridization and washing temperatures (48°C and 50°C respectively) to increase further stringency were performed, but no improvement was obtained. The three Myco378 probes were more specific using 45%-50% formamide concentration in general. Fluorescence on *Campylobacteria* cells remained weak and rare for Myco378-1 (Supplementary Fig. 4, Supplementary Fig. 5, Supplementary Table 2), looking clearly as non-specific hybridization signal (as entire *Campylobacteria* cells hybridized clearly with Epsy-549 probe). For Myco378-2 and more particularly for Myco378-3, few segments of *Campylobacteria* gave a fluorescent signal for 45% and 50% formamide.

Thus, the optimal conditions retained to ensure the best specificity of these probes is a hybridization at 46°C for 3 hours with a buffer hybridization containing 45% formamide, washing at 48°C for 30 minutes (Supplementary Table 2). The probe Myco378-3 being the less specific (a signal still being observed on *Campylobacteria* cells with this probe whatever the conditions of use), it was not further used for experiments. The probe Myco378-1 showing the best results in terms of signal specificity and efficiency was further used in the different FISH experiments on foreguts.

### Distribution of Mycoplasmatales and Deferribacteres in the digestive tract of Rimicaris spp

#### Deferribacteres in the midgut tube of R. exoculata and R. chacei

Based on previous DNA sequencing data, *Deferribacteres* were expected to be localized in the midgut of *R. exoculata* and *R. chacei* (Durand et al., 2009, 2015; Apremont et al., 2018). The Def1229 probe, specific to *Deferribacteres* digestive symbiont lineage, was used to explore the location of these bacteria in the shrimps. FISH observations of the midgut tube sections allowed us to validate the presence of these bacteria and to identify them clearly. They appeared as those previously observed with FISH using the general Eub338 probe (Durand et al., 2009) (Fig. 4A). *Rimicaris* sp. *Deferribacteres* are the long and thin cells, inserted between the microvilli of the midgut tube epithelial cells and visible in the lumen of the midgut tube, between the intestinal epithelium and the peritrophic membrane (Fig. 4A-C). They are present all along the midgut tube (Fig. 4A-C). These bacteria are easily recognizable from *Gammaproteobacteria* or *Campylobacteria* as they form “spaghetti-like” morphologies (Fig. 4A, C). Indeed, they appear as long and thin single cells, not formed of subunits like the two other phyla mentioned above. In most observations, the bacteria were so long that they curled up on themselves, forming massive clusters on the wall cells of the midgut tube (Fig. 4B, C).

**Figure 4:**
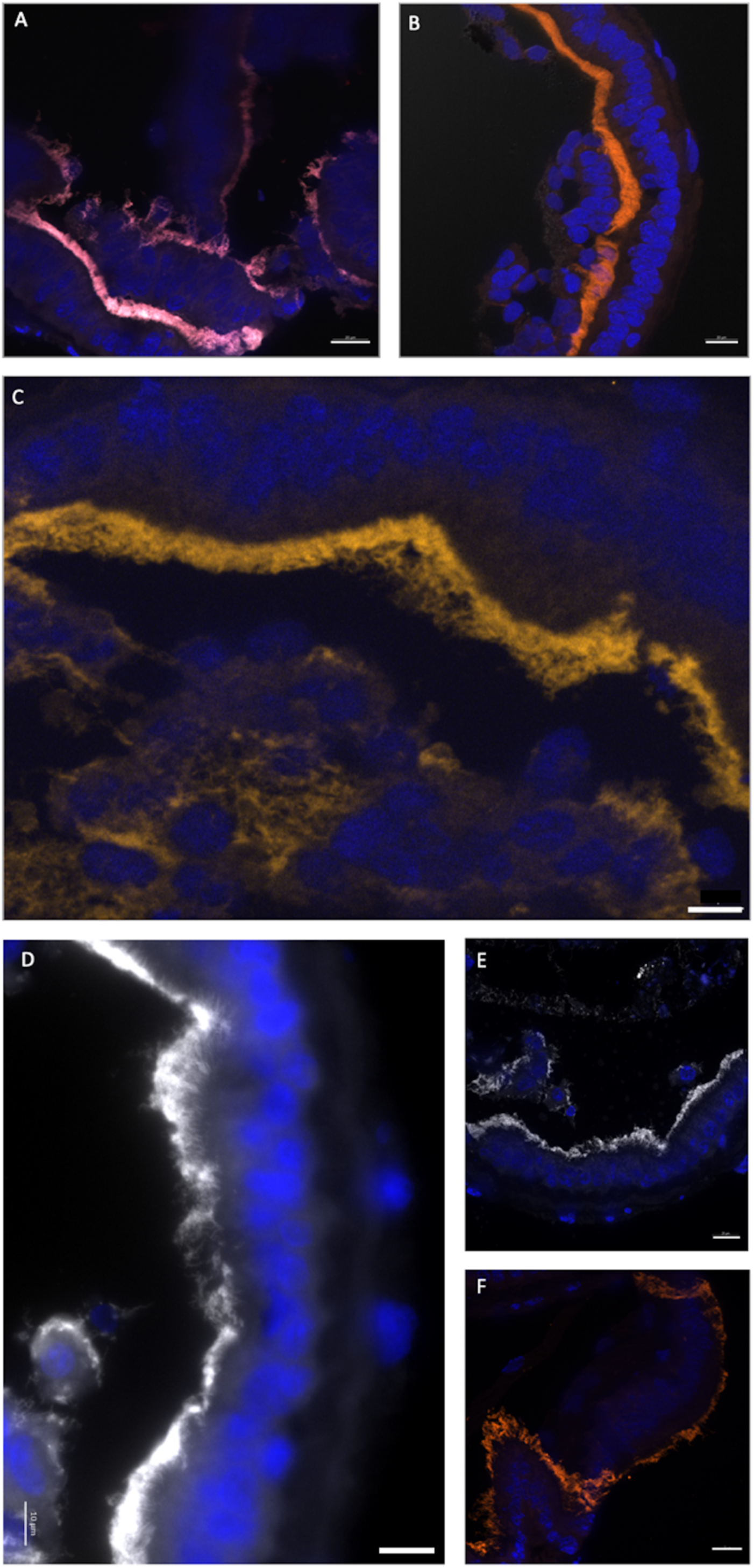
Observation of Deferribacteres in different midgut tube sections of a R. exoculata and a R. chacei adults. **(A-C)** represent sections through the midgut tube of R. exoculata adult hybridized with Eub338-cy5 (red) / Def1229-cy3 (white) **(A)** where double hybridization appeared in pink, or with Def1229-cy3 (orange in **(B)** and yellow-brown in **(C)**). Tissue cell nuclei are labeled with DAPI (blue). Formamide concentration in hybridization buffer was 30%. Pictures in **(A-C)** were taken with Apotome, and with Z-stack and DIC for B only. **(D-F)** represent sections through the midgut tube of R. chacei adult hybridized with Def1229-cy3 (white in **(D, E)** and orange in **(F)**). Tissue cell nuclei are labeled with DAPI (blue). Formamide concentration in hybridization buffer was 30%. Pictures **(E, F)** were taken with Apotome. Scale bars **(A, B, C, E, F)** = 20 µm; Scale bar **(D)** = 10 µm.

FISH observations revealed no *Deferribacteres* in the alimentary bolus, which was rich in minerals and pieces of cuticle - probably from an ingested molt. In fact, the only bacteria (few rods and coccoids) visible in the bolus hybridized only with Epsy549 and GAM42a probes. Upon dissection, the midgut tube appeared more or less transparent (i.e. more or less empty) and sometimes the alimentary bolus spilled out of the gut. FISH observations confirmed that when the alimentary bolus was absent (transparent midgut tube), *Deferribacteres* were still observable while rods and coccoids from the bolus were not. These *Deferribacteres* did not wash out with the minerals, supporting their attachment to the wall cells of the midgut. No *Deferribacteres* signal was ever revealed on the scaphognathites nor in the foregut sections, confirming their strict location to the midgut tube.

*R. chacei* midgut tube sections from two specimens were also hybridized with the Def1229 probe and results were comparable to that of *R. exoculata* (Fig. 4D-F). As for *R. exoculata*, the cephalothoracic cavity and foregut were deprived of *Deferribacteres*, contrasting with the midgut tube (Fig. 4D-F). *Campylobacteria* and *Gammaproteobacteria* cells were present only in the alimentary bolus of the midgut tube, showing coccoid and bacilli shapes. Minerals were less numerous in the alimentary bolus of both *R.* chacei *specimens*. Instead, it was filled with pieces of cuticle, more present than for *R. exoculata*. Of note, the *Deferribacteres* seemed longer on both *R. chacei* specimens, compared to *R. exoculata* ones.

According to our observations on *Rimicaris spp*. adult specimens, *Deferribacteres* symbionts are located in the midgut ectoperitrophic space, inserted between host epithelial cells microvilli.

#### Foregut structure description

As for omnivorous crustaceans, the digestive system of *R. exoculata* and *R. chacei* is composed of a mouth, an œsophagus connected to a foregut leading to a midgut tube that extends to the shrimp’s tail in the hindgut (excretion part) (Vogt, 2021). All around the foregut is the hepatopancreas appearing as an amorphous and oily gland, cream/orange colored. In our observations, the digestive system differed in size between our two species. In *R. exoculata*, the midgut tube was very long, while the foregut was not voluminous and quite flexible. The œsophagus was very short. The stomach appeared as a single cavity that could be mostly black-colored when containing minerals. In contrast, *R. chacei* adults had a more voluminous foregut, more rigid, closer to what is described for other shrimps (Vogt, 2021) and appearing mostly brown. The midgut tube of *R. chacei* was also relatively shorter. To date, the foregut of these two species have only been partially described (Segonzac et al., 1993; Apremont et al., 2018), requiring a more detailed analysis to better understand niche colonization by microorganisms. Due to sample limitation, a complete description is given for *R. exoculata* adults only (Fig. 5), by analogy to previous crustacean description (Lima et al., 2016; Vogt, 2021):

A) **<u>Global structure</u>** (Fig. 5A): The foregut of adults of *R. exoculata* is located dorsally in the cephalothorax (Fig. 1F). Like that of *Macrobrachium carcinus* (Lima et al., 2016), the overall structure seems reduced to a single cavity (Fig. 5B). The dorsal side is particularly bulging, while the ventral side is convex lengthwise. The anterior part of the stomach is connected to the œsophagus. Posteriorly, the stomach connects to the midgut tube, which dips ventrally slightly before rising to reach the dorsal side of the abdomen. The foregut is sometimes difficult to locate because of its small size hidden in the glandular mass of the hepatopancreas. The hepatopancreas is connected to the stomach ventrally. It initially surrounds the posterior ventral part of the stomach and ascends towards the dorsal face.
B) **<u>Oesophagus</u>** (Fig. 5A): This structure is located upstream of the stomach. It is a short,curved and overall narrow structure, which dips slightly from the anterior part of the stomach to the shrimp’s mouth (Fig. 5C-D). The œsophagus is lined with a thin cuticle and is surrounded with muscles. The internal cuticle of the ventral side is covered with numerous long, thin setae (here called “needles”). These are probably filtering structures that allow an initial selection of the material/minerals swallowed by the shrimp, as described for other species (Vogt, 2021).

**Figure 5:**
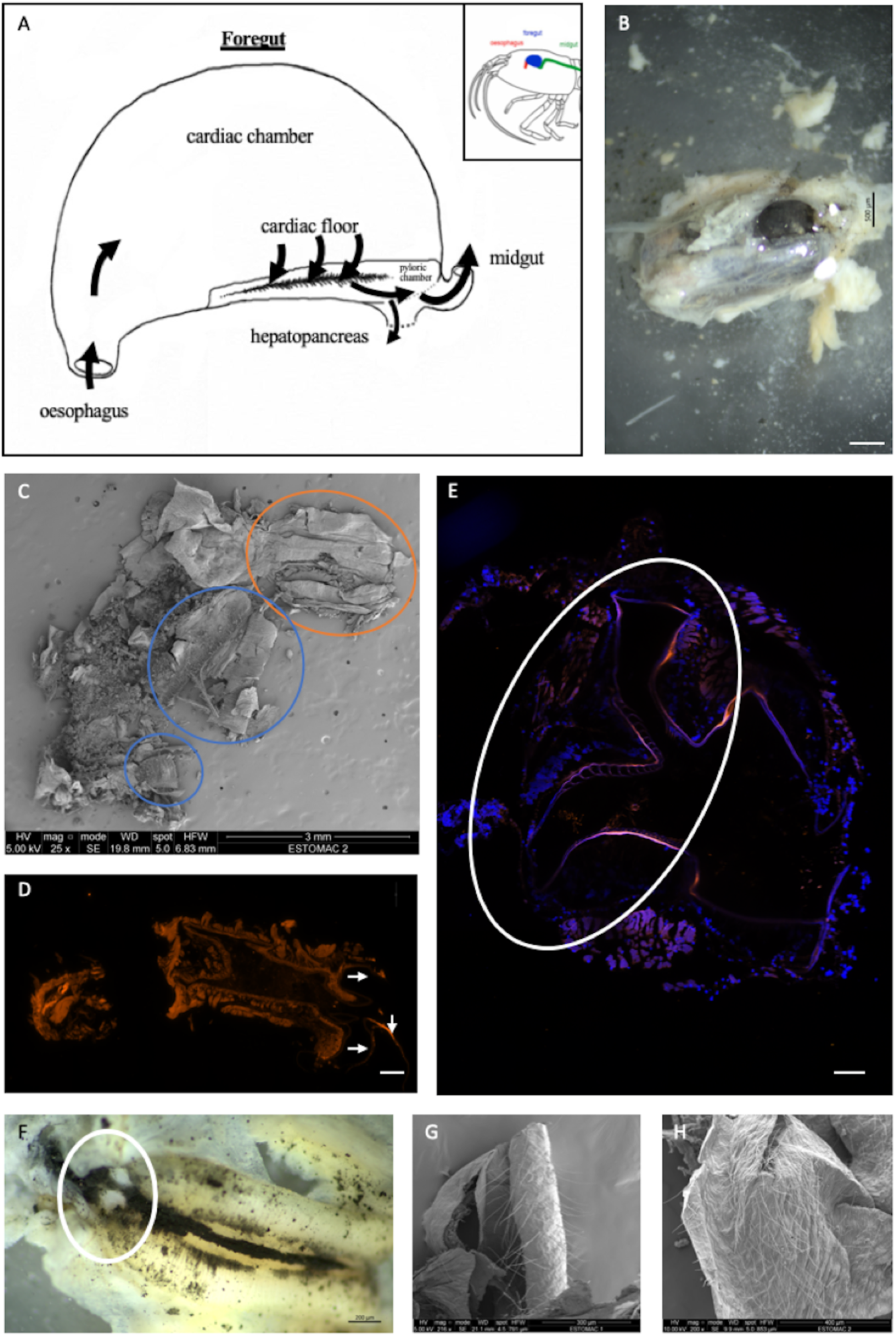
General anatomy of the foregut of a R. exoculata adult. **(A)** Drawing of the general shape of a foregut of R. exoculata adult (see Fig. 1F). The black arrows indicate the path of the ingested materials through the stomach. **(B)** Stomach (black colored) dissected from the cephalothorax surrounded by hepatopancreas (white colored) of R. exoculata (binocular microscope, scale bar = 500 µm). **(C)** Opened critical-point dried foregut revealing its content (SEM, scale bar = 3 mm). The orange circle shows the œsophagus and the blue circles show the cardiac floor sieve which is broken in two parts. **(D)** Photography of the transversal section of an œsophagus observed by autofluorescence in orange with 548 wavelength (image mosaic from 5734 x 11264 µm sections, scale bar = 20 µm). The white arrows show the ossicles where are located the setae.**(E)** Transversal section of the pyloric chamber observed by autofluorescence in pink with 548 wavelength, image mosaic from 7577 x 7577 µm sections, blue DAPI staining showing host cell nucleus (image mosaic from DIC, Apotome, scale bar = 20 µm). The circle encloses the different plates and setae – view in transversal section-present in the pyloric chamber. **(F)** Cardiac floor sieve, with the different setae and minerals (binocular microscope, scale bar = 200 µm). The circle corresponds to the probable reduced pyloric chamber. **(F-G)** Inner part of the cardiac chamber showing thin and long setae (SEM). Scale bar = 300 µm (**E**); Scale bar = 400 µm **(F)**.
C) **<u>Internal structure</u>** (Fig. 5A): The stomach is composed of two chambers - looking mostly like those of the shrimp *M. carcinus* - called the “cardiac sac or chamber”, and the “pyloric chamber” (Fig. 5C-F). The cardiac chamber is the larger of the two. It extends laterally towards the posterior part of the stomach and covers the entire pyloric chamber. The latter, which is much less voluminous and hardly visible, is located behind the cardiac floor of the cardiac chamber, in the curved ventral zone of the stomach.
D) **<u>Cardiac chamber</u>** (Fig. 5A): This is a “simple” and particularly large structure of the stomach of *R. exoculata*, which actually comprises most of the volume of the stomach (Fig. 5B). Its walls are made of striated muscles lined with a cuticle looking thinner than usually observed in other shrimps. This cuticle is covered with spicules and long thin setae (Fig. 5G-H). This chamber contains the shrimp’s alimentary bolus. Food items that come directly from the œsophagus are initially stored in this chamber (Fig. 5C). The cardiac chamber contracts and relaxes thanks to its striated muscles in movements similar to the beating of a heart. Thus, the content of the alimentary bolus is pushed toward the cardiac floor where the cardio-pyloric valve is located, and then further to the pyloric chamber. The setae, observed covering the cuticle of the cardiac chamber, provide additional filtration (Fig. 5G-H).
E) **<u>Cardiac floor</u>** (Fig. 5A): The cardiac floor is the ventral part of the cardiac chamber. It is easily recognizable because of its “bulb” shape when viewed dorsally (Fig. 5F). This floor is made of a combination of ossicles and setae, which give it a curved and thick shape, forming the cardiac floor sieve. The different structures that form the cardiac floor (according to the nomenclature of Lima et al., 2016) are the following (Fig. 6) (starting from the outermost to the innermost part of the structure):

**Figure 6:**
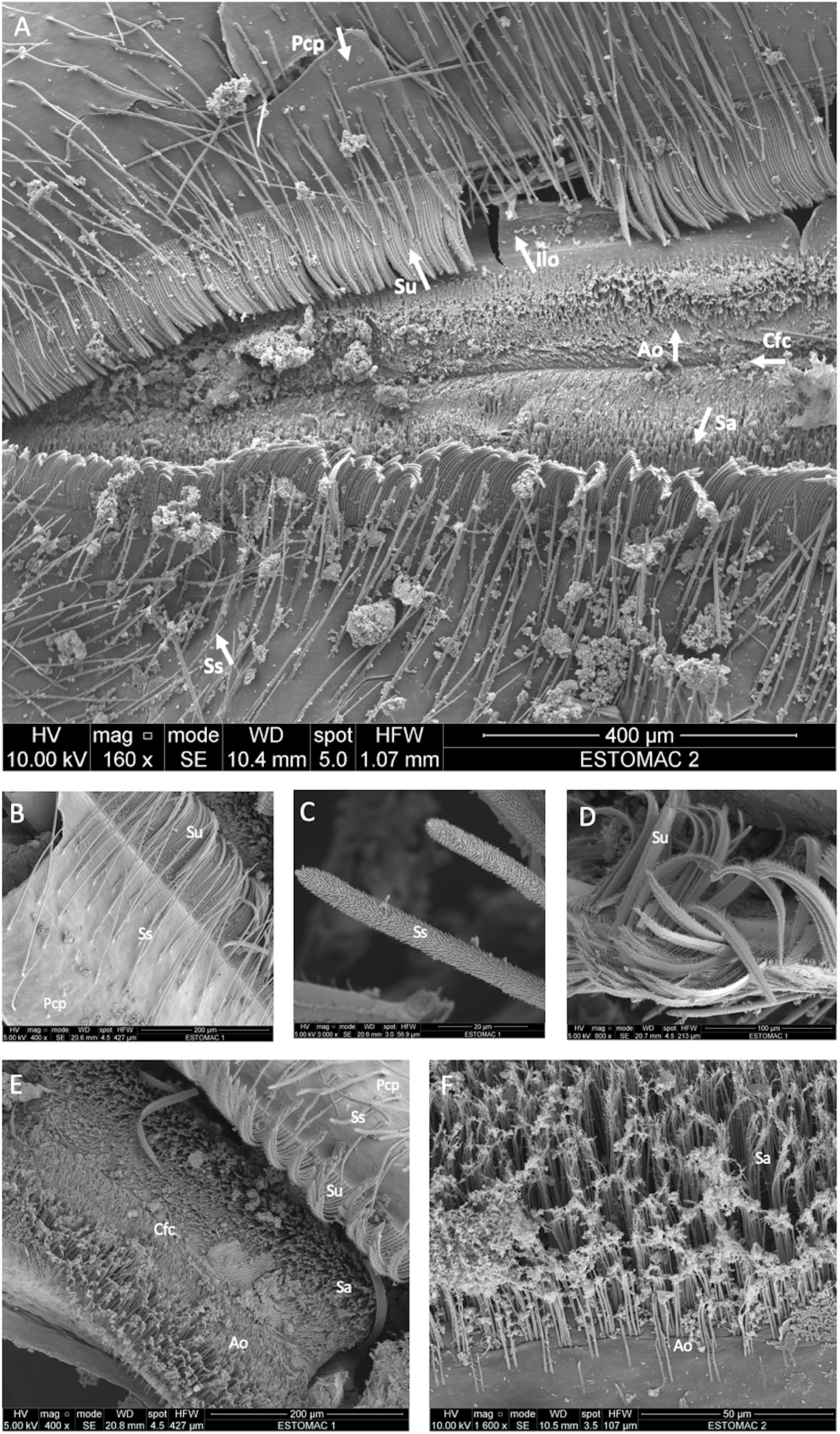
Structure of the cardiac floor of a R. exoculata adult observed with scanning electronic microscopy. **(A)** Dorsal view of the cardiac floor sieve showing the different ossicles and setae (scale bar = 400 µm). **(B)** The posterior cardiac plate with simple and serrulate setae (scale bar = 200 µm). **(C)** Simple setae of the posterior cardiac plate (scale bar = 20 µm). **(D)** Serrulate setae of the posterior cardiac plate (scale bar = 100 µm). **(E)** Closer view of the anterior ossicle with its setae, minerals spread on the surface and the cardiac floor crest on the center (scale bar = 200 µm). **(F)** The different setae of the anterior ossicle (scale bar = 50 µm.) Pcp = Posterior cardiac plate. Ss = Simple setae. Su = Serrulate setae. Ilo = Inferior lateral ossicle. Ao = Anterior ossicle. Sa = Setae of the anterior ossicle with minerals. Cfc = cardiac floor crest. - A pair of posterior cardiac plates (Fig. 6A-B). On these two plates, long setae are visible. These are simple, thick setae, but not dense (Fig. 6A-C, Fig. 7C). Along the central edge of these two plates, numerous serrulate setae (Fig. 6A-B, Fig. 7C) form a thick barrier (called sieve) between the internal structure of the cardiac floor and the two plates. These setae are thick, curved and are themselves covered with small, thin setae (Fig. 6C). By comparison, these serrulate setae remind that of the setae of the scaphognathites and exopodites of *R. exoculata* (Fig. 6C). These setae cover the paired inferior lateral ossicles (Fig. 6A, E) identified as the cardio-pyloric valve (Lima et al., 2016), and may have a role in filtering and transporting resources to the midgut (hepatopancreas and midgut tube).

**Figure 7:**
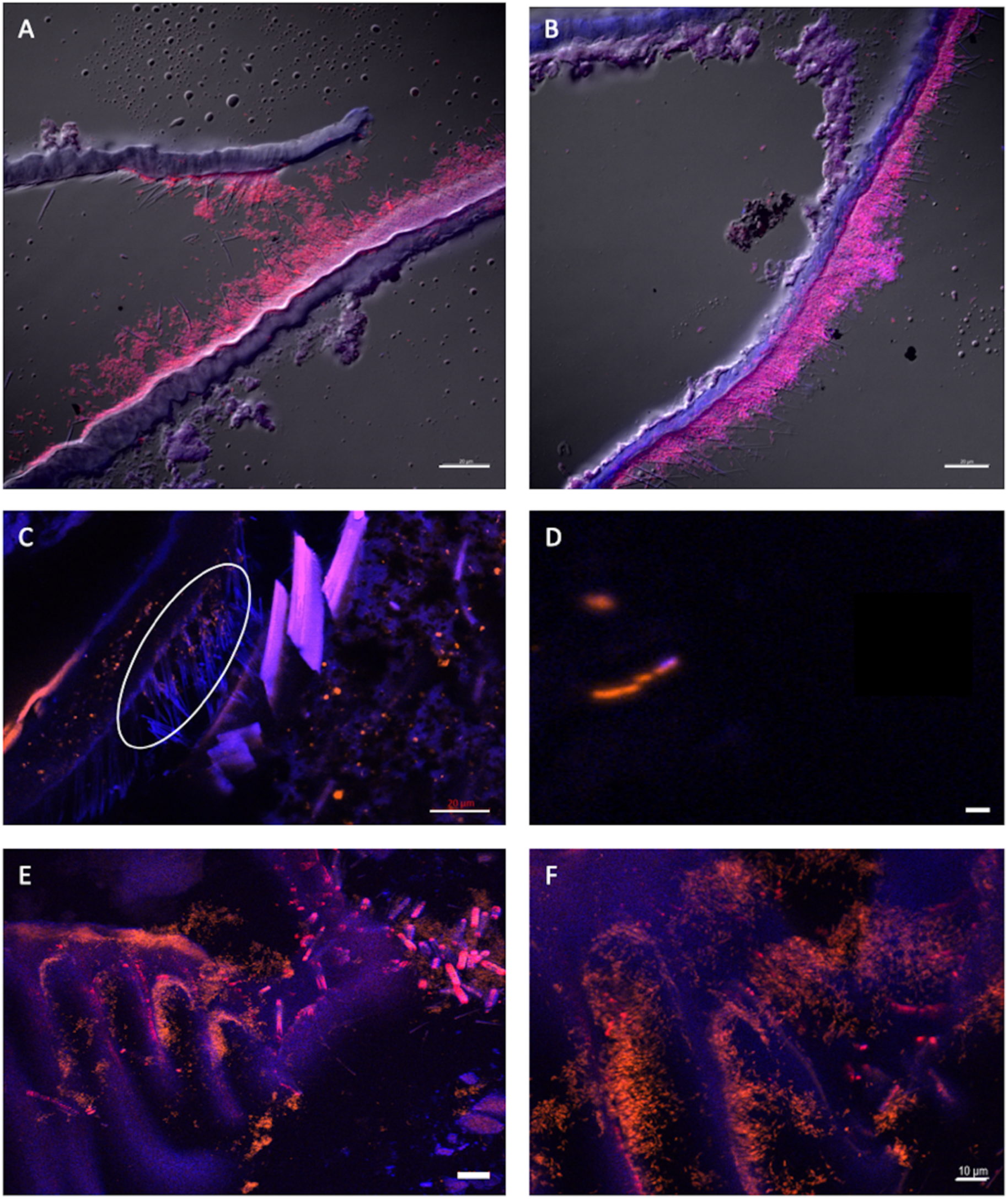
Presence of Mycoplasmatales in the foregut of R. exoculata and R. chacei adults. **(A-D)** represent sections through the foregut (œsophagus for **(A, B)**, pyloric chamber for **(C, D)**) of a R. exoculata adult hybridized with Myco378-1-cy3 (pink in **(A, B)** or orange in **(C,D)**). Tissue cell nuclei are labeled with DAPI (blue). Formamide concentration in hybridization buffer was 45% **(A-D)**. Pictures **(A-D)** were taken with Apotome, and DIC for **(A, B)**. The white circle **(C)** highlights the Mycoplasmatales hidden by the setae. **(E-F)** represent sections through the foregut (pyloric chamber) of a R. chacei adult hybridized with Myco378-1-cy3 (orange). Tissue cell nuclei are labeled with DAPI (blue). Formamide concentration in hybridization buffer was 45%. Pictures **(E, F)** were taken with Apotome. Scale bars = 20 µm **(A, B, C, E)**; Scale bar = 1 µm **(D)**; Scale bar = 10 µm **(F)**. - The unpaired anterior ossicle (of the cardio-pyloric valve (Lima et al., 2016)) is located between the pair of inferior lateral ossicles, and is visible between the serrulate setae of the posterior cardiac plates. This structure is a thin triangular calcified plate, covered with simple setae (Fig. 6A, E, F). These setae are very thin and much shorter than those found on posterior cardiac plates. They are covered with the minerals and materials contained in the shrimp’s alimentary bolus showing a filtration role, which sometimes make them difficult to distinguish (Fig. 6F). - In the center of the anterior ossicle, a longitudinal groove is visible (Fig. 6A, E). This is the cardiac floor crest, in which the residues of the alimentary bolus accumulate. The residues are brought to the pyloric chamber through this groove. The various and numerous setae allow the passage of nutrient and mineral residues to the pyloric chamber and then to the midgut (hepatopancreas and midgut tube).
F) **<u>Pyloric chamber</u>** (Fig. 5A, Supplementary Fig. 6): This structure is considerably reduced and hardly visible in *R. exoculata* adult stomach. It is a very small chamber that connects the cardiac floor to the midgut tube and is probably also connected to the hepatopancreas (Fig. 5A, 5E, 5F), but this last connection was not properly observed here due to sample limitation. By analogy to what is described for omnivorous shrimps, and according to anatomical similarities, the cardiac floor sieve separates the cardiac sac and the pyloric chamber. The cuticle of the pyloric chamber appears to be very curved and covered with dense setae that are folded on themselves (Fig. 5F, Supplementary Fig. 6). These setae are probably involved in filtration activity (see Štrus et al., 2019).

The structure of the foregut of *R. exoculata* strongly resembles that of *M. carcinus* shrimp (Lima et al., 2016) but is much smaller. In comparison, the foregut of the single available *R. chacei* specimen observed under the binocular microscope is larger than that of *R. exoculata*, with a more developed pyloric chamber, but full description is needed with new specimens.

### Mycoplasmatales in the foregut of R. exoculata and R. chacei

*Mycoplasmatales* are wall-less bacterial cells. It is extremely complicated to determine any morphology as it may change from rod to coccoid shapes. The use of the Myco378-1 probe revealed for the first time their morphology in both *Rimicaris* spp. with FISH approaches (Fig. 7A-F). To improve symbionts description in terms of both morphology and distribution on host tissues, many foregut sections were used all along the foregut from the œsophagus toward its posterior part. Depending on foregut samples, section location in the organ, fixation quality (directly on board or fixed at the laboratory after freezing), or molt stage (cuticle being more or less present and thick), the *Mycoplasmatales* were not always clearly observable (Fig. 7C).

Here for both holobionts, *Mycoplasmatales* were observed as rods only. Many divided rod-cells were observed, suggesting cell division and therefore an active community (Fig. 7D). In the different specimens studied, symbionts were observed in several areas of the foregut. First, many *Mycoplasmatales* were found in the œsophagus (Fig. 7A-B). Indeed, these symbionts accumulated as thick mats along the thin setae. The same rod morphologies were observed in the pyloric chamber in dense colonies for both *Rimicaris* spp. (Fig. 7C, E, F). Of note, the *Mycoplasmatales* colonized much more densely the two *R. chacei* specimens, mostly in the pyloric chamber on setae (Fig. 7E-F). In contrast, no *Mycoplasmatales* was observed in the food bolus, nor along the walls of the cardiac sac (Fig. 7C, E, F). Furthermore, none was evidenced at the entrance to the midgut tube. A single available *R. exoculata* adult empty foregut (i.e. without minerals) could be observed. In this case, *Mycoplasmatales* were visible on the setae of the pyloric chamber and in the lumen of the alimentary bolus. Myco378-1 gave no positive signal on any of the midgut tube sections of an adult *R. exoculata*. No fluorescence hybridization signal was ever observed on “spaghetti-like” filaments nor in the alimentary bolus (Supplementary Table 2). The *Mycoplasmatales* were thus visible along the foregut only. They were mainly associated to individual setae of the œsophagus or pyloric chamber (Fig. 7), and exhibited a rod shape, some showing evidence of active division.

## Discussion

### Rimicaris spp. midgut tube houses a tight and specific symbiosis

Unaffiliated single-cell bacteria were previously observed using transmission electron microcopy (TEM) and revealed by FISH with the universal probe Eub338 in the midgut tube of *Rimicaris exoculata* adults (Durand et al., 2009, 2015). These preliminary tests confirmed the presence of bacteria in the midgut tube, but did not allow their identification. The experiments carried out in the present study have highlighted for the first time the main lineage of the midgut bacterial community with a specific probe targeting *Deferribacteres* in both *R. exoculata and R. chacei*. *Deferribacteres* colonize all along the midgut, even for specimens having been subjected to a long fasting ((Durand et al., 2009); courtesy of Durand slides). The microorganisms are inserted between the microvilli of the midgut epithelium cells and are thus separated from the food bolus by the peritrophic membrane. Since they are located out of the alimentary bolus, *Deferribacteres* may not participate to the digestive process of the shrimp (Durand et al., 2009; Apremont et al., 2018; Zbinden and Cambon-Bonavita, 2020). Unlike the foregut or cephalothoracic cavity, there is no exuviation in the midgut tube, being of endodermic origin (Štrus et al., 2019). Thus, it is not subject to bacterial turnover with (re)colonization at each molt every 10 days (Corbari et al., 2008). The colonization of the midgut tube by *Deferribacteres* is a long-term duration association along the shrimp life cycle. Sequencing data obtained after a 72-hour fasting in adults (Durand et al., 2009) as well as our FISH observations on full or empty midgut tubes confirmed this. Indeed, whatever the nutritional state of the host, *Deferribacteres* were attached to the microvilli of the epithelium cell and can be confirmed as resident.

Filamentous bacteria are usually made of several subunits resulting from cell division, as observed for *Campylobacteria* or *Gammaproteobacteria* lineages in the cephalothoracic cavity of *Rimicaris* shrimps. TEM observations showed a different morphology for *Deferribacteres* observed in the midgut tube of *R. exoculata* (Durand et al., 2009; Apremont et al., 2018). These microorganisms do not undergo cell division (Durand et al., 2009; Zbinden and Cambon-Bonavita, 2020), although they appear highly active according to FISH results. FISH procedure being based on probe hybridization of rRNA molecules located in the ribosomes, inactive or poorly active cells are expected to have lower ribosome content and so, would give weak or even no signal, which is not the case of these *Deferribacteres*. Our observations also suggest a significant control of the host on this active symbiont lineage (cell division and colonization). Indeed, if uncontrolled bacterial growth was taking place, the alimentary bolus may be ultimately obstructed by continuously extending bacterial filaments, which was not the case here. Our FISH results in co-hybridization with Def1229 and Eub338 probes, showed that this community is almost exclusively composed of *Deferribacteres.* This further suggests a control by the host, and/or tight recognition mechanisms preventing infestation by other lineages.

Our FISH observations suggest visual differences between *Deferribacteres* in our two-host species: they appear as longer cells in both observed *R. chacei* specimens. However, since a low number of individuals were examined here, the variation in *Deferribacteres* cell length may not be attributable to the host species only, but also to intraspecific variability related to the host life stage. Additional observations including a larger set of specimens are necessary to evaluate the respective influence of host diet (mixotrophic in *R. chacei* vs symbiotrophic in *R. exoculata*), life stage, molt stage, individual nutritional status, etc…on the *Deferribacteres* lineages within the midgut tube.

According to our observations, *Deferribacteres* cells are in direct contact with their host. This contradicts previous reports on shrimps and crustaceans more generally (Martin et al., 2020; Vogt, 2021). In *Artemia salina*, *Daphnia magna*, *Gammarus* sp., *Homarus americanus*, *Tigriopus californicus* and *Sicyonia ingentis*, microorganisms appear in the midgut tube and hindgut of crustaceans, located in the alimentary bolus within the endoperitrophic space (i.e. the alimentary bolus). The peritrophic membrane that separates the endoperitrophic space and the ectoperitrophic space (i.e. between the peritrophic membrane and the epithelial cell microvilli) usually protects the epithelium against pathogens and more generally prevent any bacterial colonization (Martin et al., 2020). For the six above-mentioned species, microorganisms are ingested during feeding and washed out with the feces (considered as not essential for nutrition and digestion of their host). Our results show that *Deferribacteres* are located between the microvilli of the epithelium, within the ectoperitrophic zone, which is not “sterile” in these *Rimicaris* species, contrasting with observations made in many other crustaceans (Martin et al., 2020). This exceptional localization strengthen the hypothesis that the *Deferribacteres* play a specific role in the holobiont.

However, this role in the midgut tube is still enigmatic and a metagenomics study recently showed that this lineage would be involved in host nutrition and defense (genes coding for fumarate reductase, genes required for biotin and riboflavin biosynthesis, immunity systems, Aubé, Cambon-Bonavita et al., submitted). Their high abundance in both *Rimicaris* spp. may provide a protection against proliferation of deleterious bacteria. This is the case of *Deferribacteres* found in mammals, particularly in rodents (Lee, 1985). In rats and mice, *Deferribacteres* relatives (new genus and species: *Mucispirillum schaedleri* (Robertson et al., 2005)) are located in the mucus layer covering and protecting the gastrointestinal epithelial cells (Lee, 1980). This mucus also harbors other spiral bacteria belonging to the genus *Helicobacter* and *Campylobacter* (Lee et al., 1968; Davis et al., 1972), as well as *Salmonella enterica serovar Typhimurium* which causes non-typhoidal colitis in hosts (Herp et al., 2019). *Deferribacteres* relatives are spiral-shaped bacteria that play a key protective role in the symbiosis of the digestive system as they outcompete pathogenic *Salmonella enterica serovar Typhimurium* (Herp et al., 2019). Similarly, in *Rimicaris* shrimps, *Deferribacteres* may act as a barrier against deleterious microorganisms thus having a double role: protection of the host through competitive interactions with other bacteria, and host nutrition. They may also play a role in detoxification of minerals contained in the alimentary bolus.

### A foregut functioning and symbiosis

In omnivorous shrimps, the foregut is composed of two chambers, covered with tooth, setae and spicules allowing food processing and filtration of nutriments. The food bolus contained in the cardiac chamber is directed towards the pyloric chamber by means of setose lateral ridges and lateral valves avoiding regurgitation. Passing through sieves, alimentary bolus smallest particles (less than 1µm) enter the hepatopancreas (a major digestive and storage gland in crustaceans) and larger ones are directed to the midgut tube, embedded in the peritrophic membrane. Thus, the stomach triturates food and act as a filter, while the hepatopancreas and midgut tube are involved in nutrient absorption and digestion (Pattarayingsakul et al., 2019; Štrus et al., 2019). Our anatomical study of the stomach of *Rimicaris* spp. also revealed complex filtering structures. In *R. exoculata*, the very small size of the foregut and its thin cuticle reinforce the idea of a weak mechanical action (Segonzac et al., 1993; Durand et al., 2009), reminding what is described for the shrimp *Macrobrachium carcinus* (Lima et al., 2016). In *M. carcinus*, the pyloric and cardiac chambers form a nearly single cavity (Lima et al., 2016), with almost no grinding appendages – no teeth in the œsophagus, nor in the cardiac chamber, like other crustaceans (Vogt, 2021). In this species, gastric mill and food trituration is possible thanks to the high quantity of sand ingested by the shrimp (Lima et al., 2016). By analogy, in *R. exoculata,* ingested minerals may play the role of the grinding appendages, which are absent in this species (no teeth) and the alimentary bolus would then be directed toward the midgut through the pyloric sieve. In *R. chacei*, the foregut is almost twice as large as that of *R. exoculata,* with a thicker cuticle. This, together with the presence of functional chelipeds (Casanova et al., 1993), numerous pieces of cuticle and organic material in the digestive tract, strengthen the mixotrophic diet hypothesis (Apremont et al., 2018, Methou et al., 2020), with a more important foregut mechanical digestive activity.

Despite these anatomical differences, both species exhibit similar bacterial colonization of their foregut, including the œsophagus and stomach, with a higher density of bacteria observed in the two *R. chacei* specimens. *Mycoplasmatales* colonize specific areas inside the foregut of both *Rimicaris* spp. We observed them in dense clusters on the setae at the entrance of the foregut (œsophagus) or on setae of the pyloric chamber for *R. exoculata*, and in high density close to the junction with the midgut tube for *R. chacei. Mycoplasmatales* were sometimes difficult to observe in the pyloric chamber of *R. exoculata*. It can be hypothesized that they were hidden by the cuticle and setae that occupy most space in the pyloric chamber (Fig. 7C), or signal observation may be impaired due to ingested minerals. Generally, *Mycoplasmatales* seemed to colonize preferentially dense setae areas, reminding cephalothorax bacterial colonization. However, additional observations on a higher number of specimens would be required to better understand the distribution of *Mycoplasmatales* within *Rimicaris* foregut, and identify common patterns or variations related to the life stage, molt stage, nutritional status or host species.

Being densely present in all specimens, *Mycoplasmatales* may play an essential role in the digestion processes of *Rimicaris* spp. Associated with the thin setae localized in the first part of the foregut (œsophagus) they may be involved in processing minerals, food items or cuticle pieces. They are also observed on setae forming sieves of the pyloric chamber through which crushed alimentary bolus passes. Their role may then be comparable to that of the Bg1 and Bg2 *Mycoplasmatales* relatives found in the foregut of the marine isopods *Bathynomus* sp. (Wang et al., 2016). These lineages are close to *’Candidatus* Hepatoplasma crinochetorum’ found in terrestrial isopods hepatopancreas (Wang et al., 2004, 2007). In the case of *Bathynomus* sp., Bg1 and Bg2 *Mycoplasmatales* are retrieved in the foregut content, and favor host development in nutrient-poor environments (Wang et al., 2016). In the same way, in terrestrial isopods, *Mycoplasmatales* are involved in host food digestion through complementation of its unbalanced diet based on leaves (Wang et al., 2004, 2007). In terrestrial isopods, digestion occurs in the hepatopancreas, after ingested material is crushed in the foregut, and the midgut is almost absent. This differs from marine isopods and was proposed to be an adaptation to terrestrial conditions (Wang et al., 2004). Then, *Mycoplasmatales* may play the same role in the hepatopancreas of terrestrial isopods and in the foregut of marine crustaceans. In deep-sea hydrothermal environments, most nutrition, but not all, is fueled by symbiotic chemosynthesis. This may also lead to unbalanced diet, to support chitin synthesis for example, as hydrothermal shrimps have a very short molt cycle duration (Corbari et al., 2008). We can hypothesize that *Mycoplasmatales* help the host to degrade chitin of the cuticle, as described for *Porcellio scaber* (Bouchon et al., 2016). This hypothesis is supported by the occurrence of genes coding for enzymes involved in chitin metabolism in the metagenomes of the *Mycoplasmatales* symbionts (Aubé, Cambon-Bonavita et al., submitted). Overall, these observations suggest that the *Mycoplasmatales* colonization is directed toward specific location on specific structures, probably related to their role and function. Neither the host nor the organ (hepatopancreas *vs* foregut) would have an impact on their colonization capacity.

### A complex symbiosis linked to the diet of both Rimicaris spp

As the cephalothoracic cavity of both *Rimicaris* spp., their digestive system differs in their morphology. According to FISH and SEM observations on several *R. exoculata* specimens, the anatomy of the foregut has been described in more detail. We confirm that the foregut is a filtering structure as for other shrimps. The œsophagus is reduced to a simple structure with filtering elements. The stomach is deprived of grinding appendage but exhibits many setae, and a sieve to filter the nutrient and the minerals. This stomach is highly reduced, but not the midgut tube nor the hepatopancreas. The digestive system of the few *R. chacei* observed (FISH, binocular microscope) showed significant morphological differences. The stomach is much more voluminous compared to that of *R. exoculata*, with a thicker cuticle, and a larger pyloric chamber. We can hypothesize a more developed internal structure, with grinding appendages, as free mandibles permit organic food ingestion for this species. The digestive system size and structure is probably related to the diet, chemosynthetic for *R. exoculata* and mixotrophic for *R. chacei*. To conclude on this potential adaptation, more observations of *R. chacei* specimens are required, including along life and molt cycles.

This study, initially based on the development of new probes for FISH observations (Def1229 targeting *Deferribacteres* and probes Myco378-1, Myco378-2 and Myco378-3 for *Mycoplasmatales*), allowed us to complement previous molecular 16S rDNA-based studies performed on the whole digestive system of *R. exoculata* and *R. chacei* adults. Both *Rimicaris* spp. show a double functional symbiosis located on the foregut setae for the *Mycoplasmatales* and attached to the midgut epithelial cells for the *Deferribacteres.* These symbionts are proposed as new lineages (Aubé, Cambon-Bonavita et al., submitted). For both hosts, the phylogeny and the morphology of each lineage colonizing the digestive system are similar. *Deferribacteres* are single long and thin filamentous cells, while *Mycoplasmatales* are rod shaped bacteria. For the two *R. chacei* specimens studied and all the *R. exoculata* specimens, *Deferribacteres* densely colonized the midgut tube on its all length. *Deferribacteres* seem to be longer in *R. chacei*, which may be related to its diet behavior and an alimentary bolus with less minerals. The distribution of *Mycoplasmatales* is more difficult to compare between the two *Rimicaris* spp. due to a lack of *R. chacei* specimens. For *R. exoculata*, the setae of the œsophagus and of the pyloric chamber are colonized with a dense symbiont population. It was also the case for the single *R. chacei* from Snake Pit. The pyloric chamber was more densely colonized, which seemed to be linked to the morphology of the thicker cuticle and numerous setae. However, the morphology of the foregut filtering structures depends probably on the molt cycle stage and the host species (denser and thicker in the mixotrophic *R. chacei*). In the same way, the symbiont colonization is subjected to molt event and so, loss of cuticle including setae. To better localize and compare the digestive symbiosis of the two species, a new study, comprising more *Rimicaris* spp. specimens along life and molt cycles is necessary.

The digestive symbiosis of both *Rimicaris* spp. shares similarities with other species such as mammals for the midgut and other terrestrial, freshwater or marine crustaceans for the foregut. These symbioses allow both the host to develop and adapt in hostile and nutrient-poor environments, and in return allows the symbionts to shelter and colonize favorable niches. Hydrothermal shrimps show commonalities with isopods in the functioning of their foregut and with mice for midgut functioning. This suggests a somewhat “universal” evolution of digestive symbiosis. The specific location of the various microorganisms in the digestive system allows us to propose hypothesis about their role and function. The *Deferribacteres* could be mainly protective for the host, while the *Mycoplasmatales* may support host development. Their function has yet to be further studied through metagenomics and *in vivo* studies.

Knowledge gaps persist regarding the transmission of *Deferribacteres* and *Mycoplasmatales* lineages along *Rimicaris* life cycle and molt events. Because the foregut is covered with cuticle, removed during exuviation, the hypothesis of horizontal transmission is often proposed for *Mycoplasmatales.* On the contrary, the control of *Deferribacteres* cell division and colonization by the host, together with the quasi absence of their free-living relatives in water surrounding shrimps (Durand et al., 2009; Hügler et al., 2010; Flores et al., 2011; Cowart et al., 2017) let suppose a specific symbionts recognition and their vertical transmission. However no symbiont was observed on the envelope of brooded eggs (Methou et al., 2019), raising questions about potential mechanisms allowing such vertical transmission for *Deferribacteres*. Another way of transmission may be proposed, as for terrestrial isopods, where juveniles eat the feces of the adults when ingesting food on litter (Wang et al., 2007). This may be the case for the *R. exoculata* where juveniles are recruited close to adults’ aggregations (Methou et al., 2022). However, *R. chacei*, juveniles are first mainly recruited in isolated patches, without adults or few (Methou et al., 2022). This may then reduce digestive symbiont acquisition efficiency, leading to lower holobiont fitness, and may be a partial explanation of the observed population collapse of *R. chacei*. Still, more studies are required especially on juveniles to confirm or not an acquisition during recruitment. Understanding the way of acquisition of these symbionts (horizontal, vertical and/or mixed transmission) would help to understand the development of both hosts and their relation to their symbionts. The study of juveniles and larvae (Guéganton et al., in prep) is then essential to answer the questions of transmission, localization and functions of these symbionts.

## Acknowledgments

We thank cruise chief scientist of BICOSE2 2018 cruise (M.A. Cambon-Bonavita) and the captain and crew of R/V *Pourquoi pas?* and Nautile for logistic assistance in collecting samples.

