## Supplementary Figures and Tables for "Anatomy and Symbiosis of the digestive system of the vent shrimps *Rimicaris exoculata* and *Rimicaris chacei* revealed through imaging approaches"

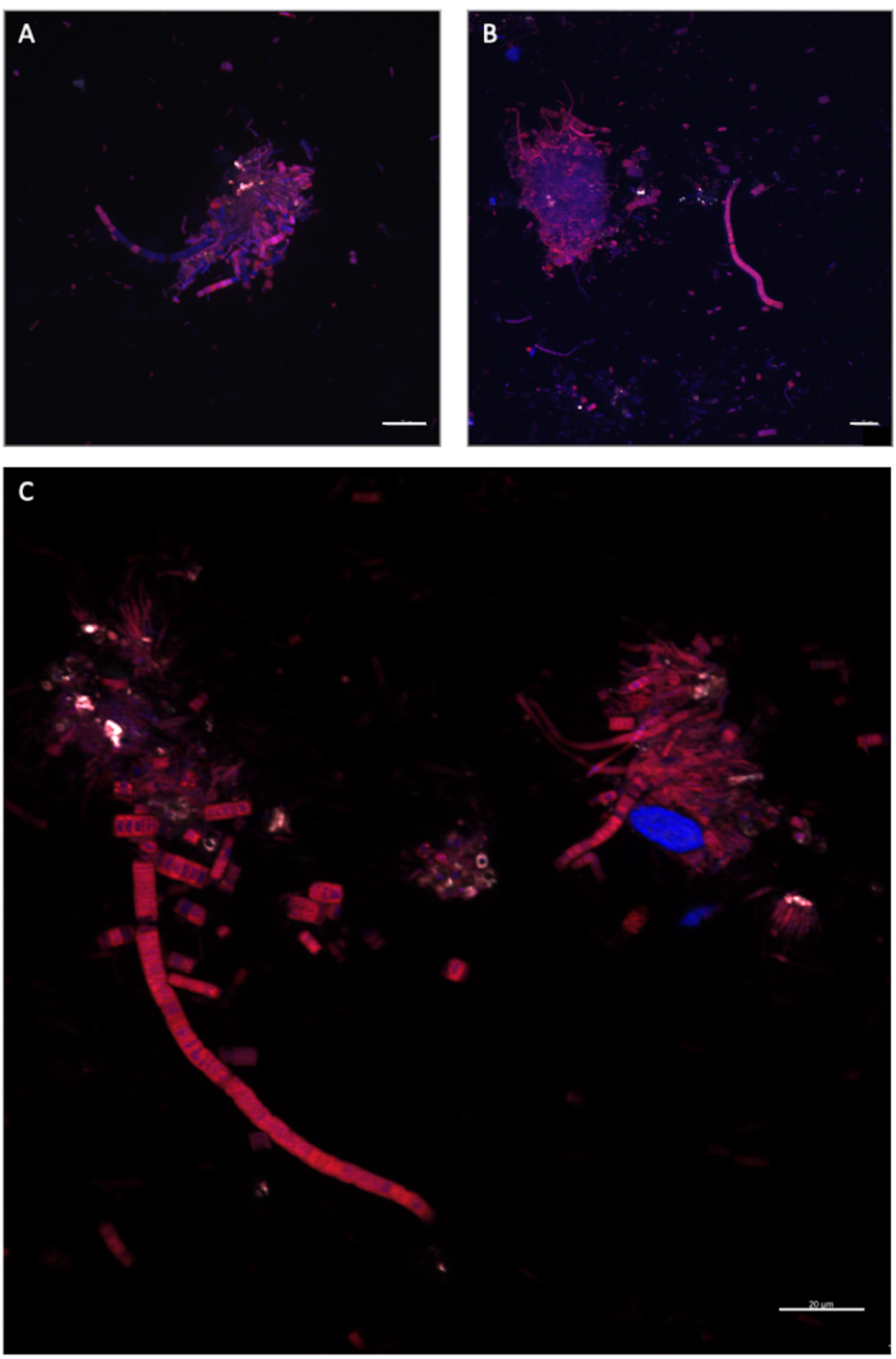


*Supplementary Figure 1: Stringency tests with Def1229/ Eub338.* ***(A-C)*** *represent sections through the scaphognathites of a R. exoculata adult hybridized with Eub338-cy5 (pink in* ***(A,B)*** *and red in* ***(C)****) /Def1229-cy3 (white)* ***(A-C)****. Tissue cell nuclei are labeled with DAPI (blue). Formamide concentration in hybridization buffer was 20%* ***(A)****, 30%* ***(C)****, 40%* ***(B)****. Pictures were taken with Apotome. Scale bars = 20 µm.*

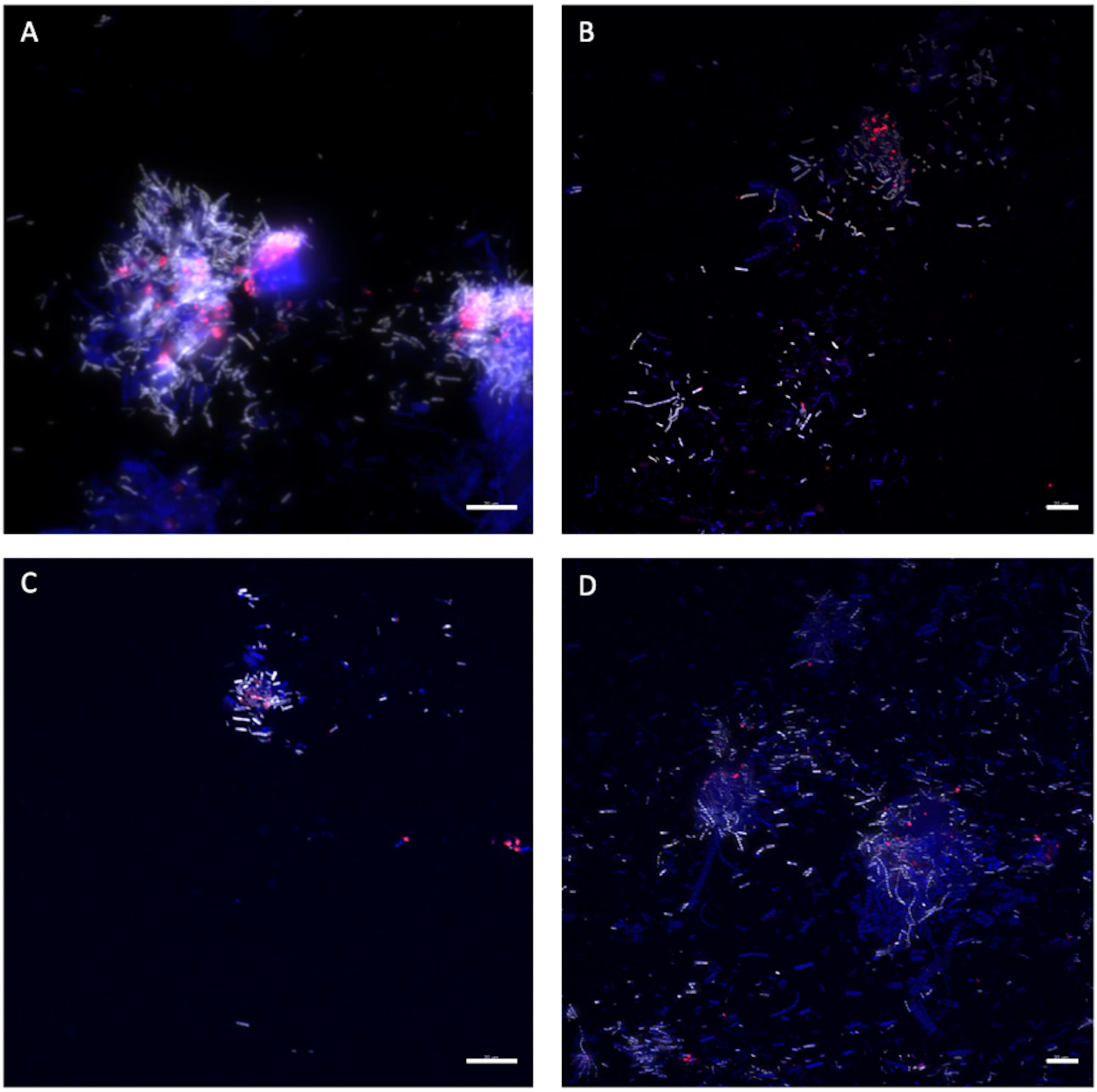


*Supplementary Figure 2 : Stringency tests with Def1229/ GAM42a.* ***(A-D)*** *represent sections through the scaphognathites of R. exoculata adult hybridized with GAM42a-cy3 (white) / Def1229-cy5 (red). Tissue cell nuclei are labeled with DAPI (blue). Formamide concentration in hybridization buffer was 20%* ***(A)****, 30%* ***(B)****, 40%* ***(C)****, 50 %* ***(D)****. Pictures* ***(B, C, D)*** *were taken with Apotome. Scale bars = 20 µm.*

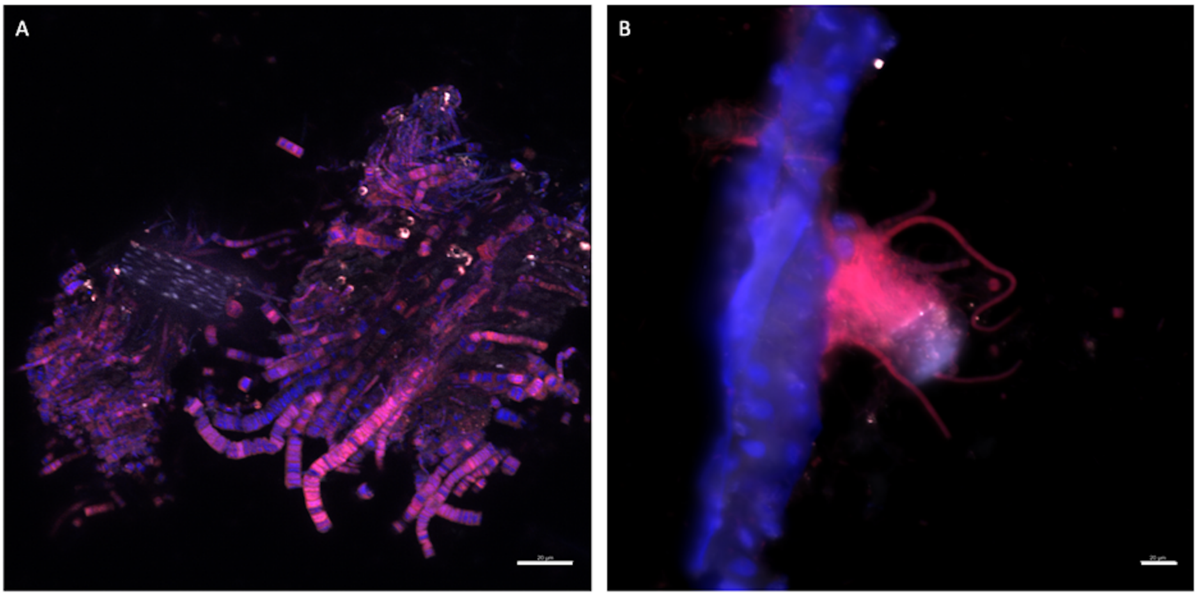


*Supplementary Figure 3: Stringency tests with Def1229/ Epsy549.* ***(A-B)*** *represent sections through the scaphognathites of a R. exoculata adult hybridized with Epsy549-cy5 (red)/ Def1229-cy3 (white).* ***(B)*** *represents a colony of Campylobacteria on host tissue. Tissue cell nuclei are labeled with DAPI (blue). Formamide concentration in hybridization buffer was 30%* ***(A)****, 40%* ***(B)****. Picture* ***(A)*** *was taken with Apotome. Scale bars = 20 µm.*

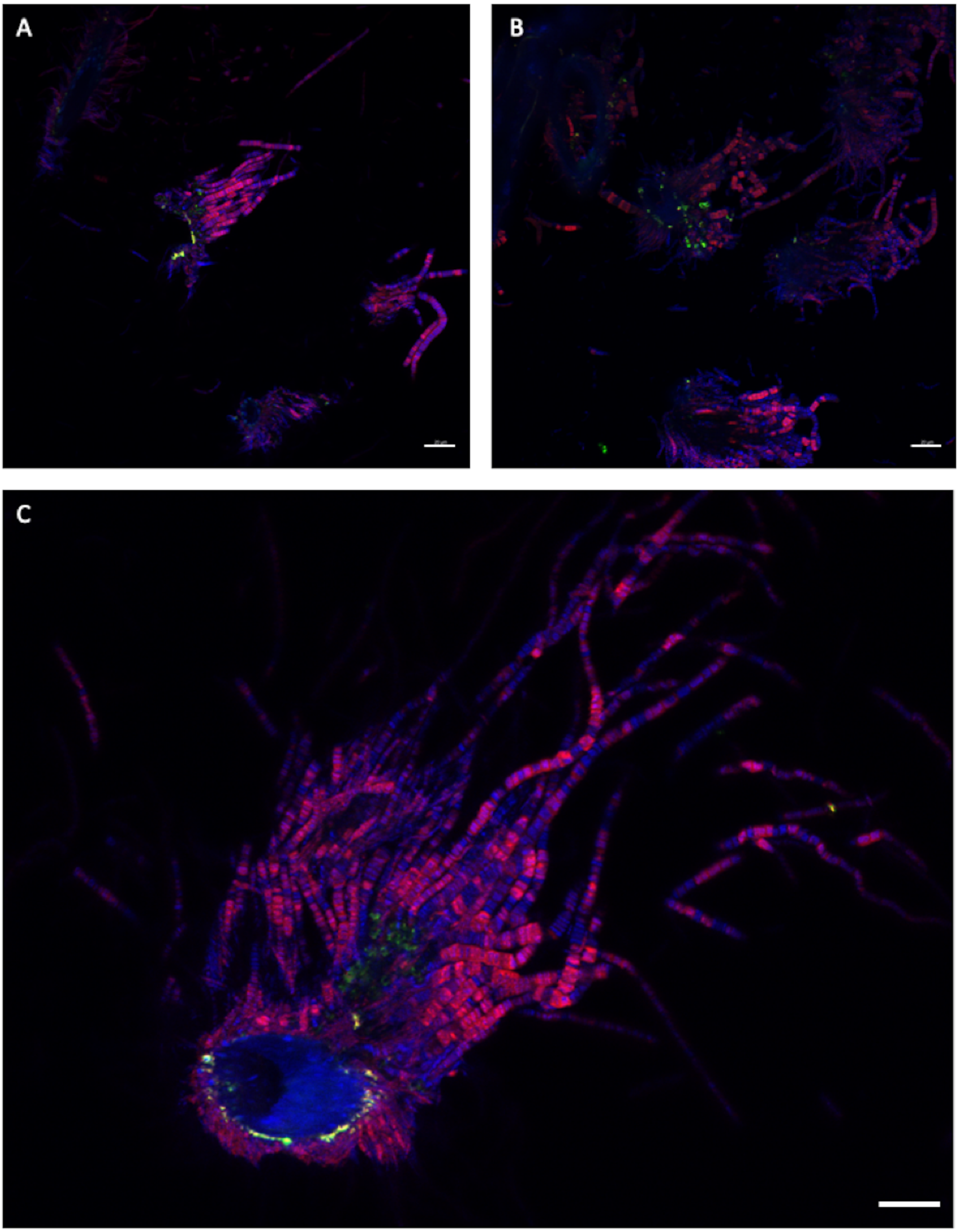


Supplementary Figure 4 : *Stringency tests with Myco378-1/Eub338, Myco3782-2/ Eub338 and Myco378-3/Eub338.* ***(A-C)*** *represent sections through the scaphognathites of R.exoculata adult hybridized with Myco378-1-cy3 (green)/ Eub338-cy5 (pink)* ***(A)*** *or with Myco378-2-cy3 (green)/ Eub338-cy5 (pink)* ***(B)*** *or with Myco378-3-cy3 (green)/ Eub338-cy5 (pink)* ***(C)****. Tissue cell nuclei are labeled with DAPI (blue). Formamide concentration in hybridization buffer was 35%* ***(A)****, 40%* ***(B)****, 50%* ***(C)****. Pictures were taken with Apotome. Scale bars = 20 µm.*

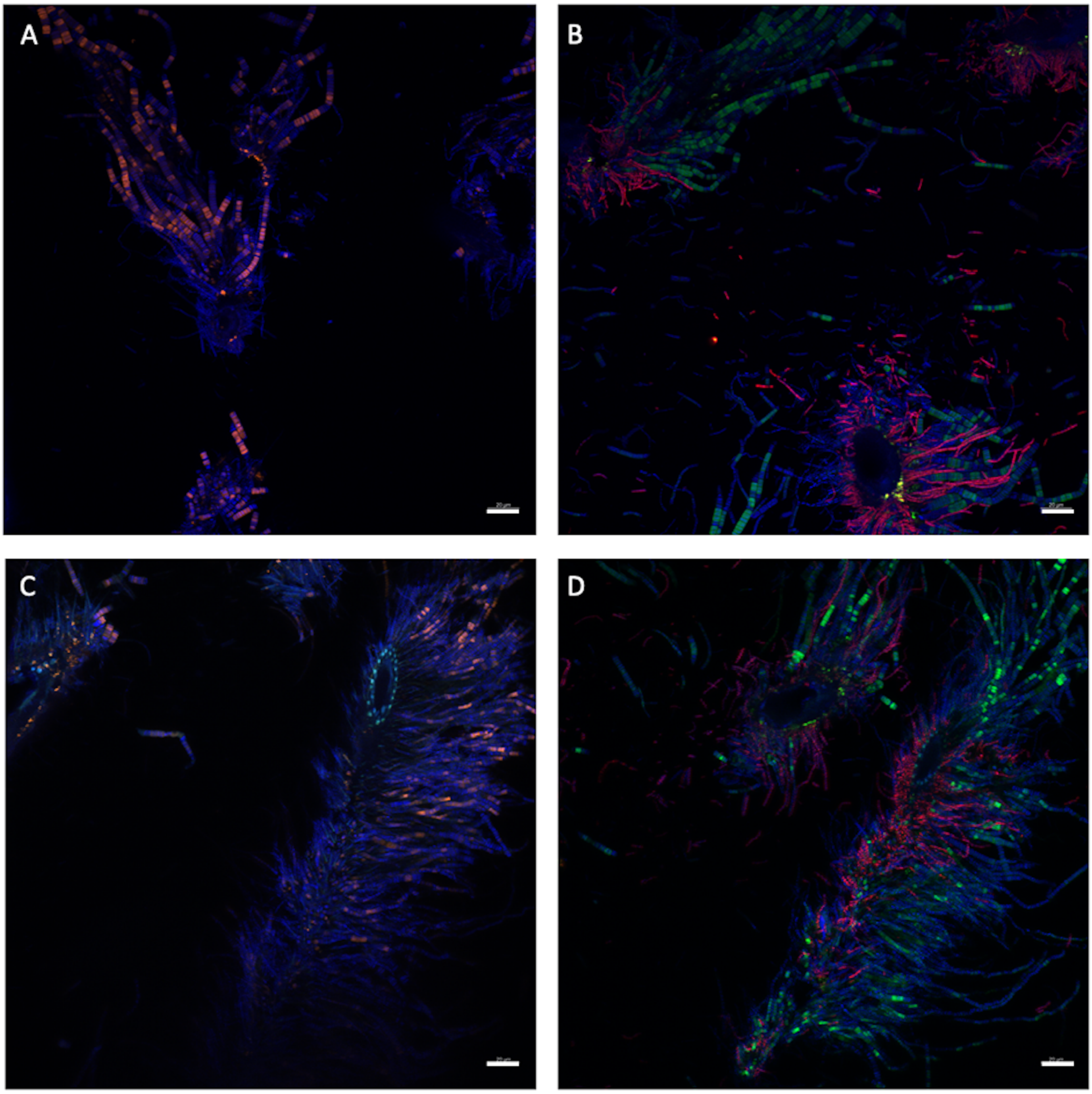


Supplementary Figure 5: *Stringency tests with Myco378-2 and Myco378-1.* ***(A-D)*** *represent sections through the scaphognathites of R. exoculata adult hybridized with Myco378-2-cy3 (orange)* ***(A)****, or with Myco378-2-cy3 (green)/ GAM42a-cy5 (red)* ***(B)****, or with Myco378-1-cy3 (orange)* ***(C)****, or with Myco378-1-cy3 (green) / GAM42a-cy5 (red)* ***(D)****. Tissue cell nuclei are labeled with DAPI (blue). Formamide concentration in hybridization buffer was 35%* ***(A,B)****, and 45%* ***(C, D)****. Pictures were taken with Apotome. Scale bars = 20 µm.*

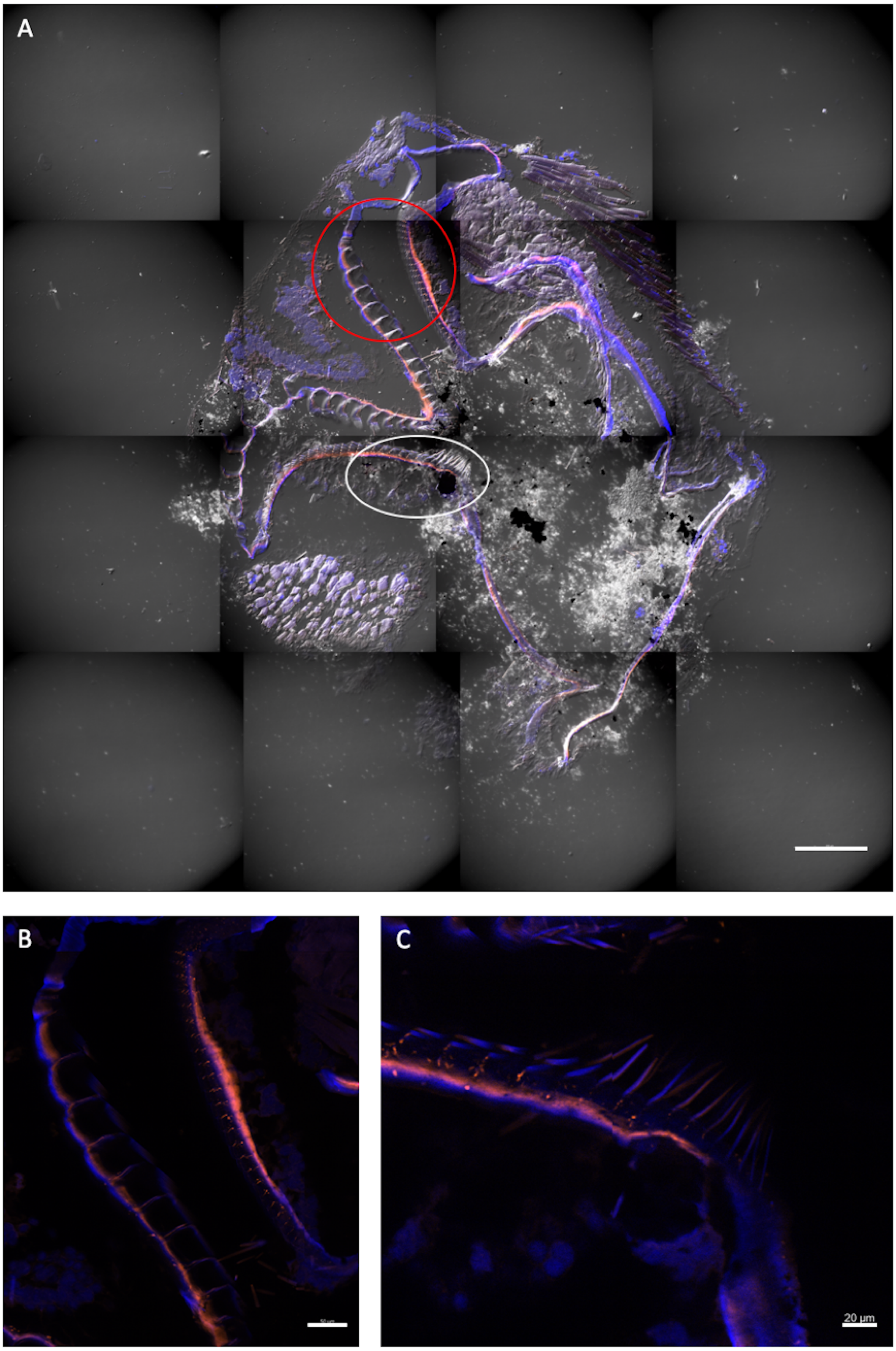


*Supplementary Figure 6 : Photography of the pyloric chamber of a R. exoculata adult in transversal section.* ***(A)*** *Entire pyloric chamber observed by autofluorescence in orange with 548 wavelength, image mosaic from 7577 x 7577 µm sections, blue DAPI staining showing host cell nucleus, DIC, Apotome, scale bar = 200 µm. The blue circle encloses the different plates and the white circle encloses the setae.* ***(B)*** *Zoom on the plates of the pyloric chamber. Scale bar = 50 µm.* ***(C)*** *Zoom on the setae of the pyloric chamber. Scale bar = 200 µm.*

### Supplementary Tables

*Supplementary Table 1 : Summary of the stringency tests performed with the probe Def1229. Probes were hybridized at 46°C, and washed at 48°C.*

| Formamide % | 20 | 30 | 40 | 50 | 20 | 30 | 40 | 50 |
|---|---|---|---|---|---|---|---|---|
| Probes\target tissue | Scaphognathite | | | | Midgut tube | | | |
| Def1229 | - | - | - | - | +++ | +++ | +++ | +++ |
| Eub338 | +++ | +++ | +++ | +++ | +++ | +++ | +++ | +++ |
| Epsy549 | +++ | +++ | +++ | +++ | - | - | - | - |
| GAM42a | +++ | +++ | +++ | +++ | - | - | - | - |
| Def1229 + Eub338 | +++ Eub338 only | +++ Eub338 only | +++ Eub338 only | +++ Eub338 only | +++ both probes | +++ both probes | +++ both probes | +++ both probes |
| Def1229 + Epsy549 | +++ Epsy549 only | +++ Epsy549 only | +++ Epsy549 only | +++ Epsy549 only | +++ Def1229 only | +++ Def1229 only | +++ Def1229 only | +++ Def1229 only |
| Def1229 + GAM42a | +++ GAM42a only | +++ GAM42a only | +++ GAM42a only | +++ GAM42a only | +++ Def1229 only | +++ Def1229 only | +++ Def1229 only | +++ Def1229 only |

*Supplementary Table 2 : Summary of the stringency tests performed with the probes Myco378-1, Myco378-2 and Myco378-3.*
** some segments of Campylobacteria gave a positive signal; ** rare segments of Campylobacteria gave a positive signal.*
*Probes were hybridized at 46°C, and washed at 48°C.*

| Formamide % | 35 | 40 | 45 | 50 | 30 | 35 | 40 | 45 |
|---|---|---|---|---|---|---|---|---|
| Probe \ target tissue | Scaphognathite | | | | Foregut | | | |
| Myco378-1 | -/+* | -* | -** | -** | ++ | ++ | +++ | +++ |
| Myco378-2 | +* | -/+* | -/+* | -/+** | ++ | ++ | +++ | +++ |
| Myco378-3 | ++* | ++* | +* | -/+* | ++ | ++ | +++ | +++ |
| Eub338 | +++ | +++ | +++ | +++ | +++ | +++ | +++ | +++ |
| Epsy549 | +++ | +++ | +++ | +++ | - | - | - | - |
| GAM42a | +++ | +++ | +++ | +++ | - | - | - | - |
| Myco378-1 + Eub338 | +++<br>Eub338 only | +++<br>Eub338 only | +++<br>Eub338 only | +++<br>Eub338 only | +<br>Myco378-1<br>+++Eub338 | +<br>Myco378-1<br>+++Eub338 | ++<br>Myco378-1<br>+++Eub338 | ++<br>Myco378-1<br>+++Eub338 |
| Myco378-1 + Epsy549 | -/+*<br>Myco378-1<br>+++Epsy549 | -*<br>Myco378-1<br>+++Epsy549 | -**<br>Myco378-1<br>+++Epsy549 | -**<br>Myco378-1<br>+++Epsy549 | ++<br>Myco378-1<br>only | ++<br>Myco378-1<br>only | +++<br>Myco378-1<br>only | +++<br>Myco378-1<br>only |
| Myco378-1 + GAM42a | -/+*<br>Myco378-<br>1+<br>++GAM42a | -*<br>Myco378-1<br>+++GAM42a | -**<br>Myco378-1<br>+++GAM42a | -**<br>Myco378-1<br>+++GAM42a | ++<br>Myco378-1<br>only | ++<br>Myco378-1<br>only | +++<br>Myco378-1<br>only | +++<br>Myco378-1<br>only |
| Myco378-2 + Eub338 | +++<br>Eub338 only | +++<br>Eub338 only | +++<br>Eub338 only | +++<br>Eub338 only | +<br>Myco378-2<br>+++Eub338 | +<br>Myco378-2<br>+++Eub338 | ++<br>Myco378-2<br>+++Eub338 | ++<br>Myco378-2<br>+++Eub338 |
| Myco378-2 + Epsy549 | +*<br>Myco378-2<br>+++Epsy549 | -/+*<br>Myco378-2<br>+++Epsy549 | -/+*<br>Myco378-2<br>+++Epsy549 | -/+**<br>Myco378-2<br>+++Epsy549 | ++<br>Myco378-2<br>only | ++<br>Myco378-2<br>only | +++<br>Myco378-2<br>only | +++<br>Myco378-2<br>only |
| Myco378-2 + GAM42a | +*<br>Myco378-2<br>+++GAM42a | -/+*<br>Myco378-2<br>+++GAM42a | -/+*<br>Myco378-2<br>+++GAM42a | -/+**<br>Myco378-2<br>+++GAM42a | ++<br>Myco378-2<br>only | ++<br>Myco378-2<br>only | +++<br>Myco378-2<br>only | +++<br>Myco378-2<br>only |
| Myco378-3 + Eub338 | +++<br>Eub338 only | +++<br>Eub338 only | +++<br>Eub338 only | +++<br>Eub338 only | +<br>Myco378-3<br>+++Eub338 | +<br>Myco378-3<br>+++Eub338 | ++<br>Myco378-3<br>+++Eub338 | ++<br>Myco378-3<br>+++Eub338 |
| Myco378-3 + Epsy549 | ++*<br>Myco378-3<br>+++Epsy549 | ++*<br>Myco378-3<br>+++Epsy549 | +*<br>Myco378-3<br>+++Epsy549 | -/+*<br>Myco378-3<br>+++Epsy549 | ++<br>Myco378-3<br>only | ++<br>Myco378-3<br>only | +++<br>Myco378-3<br>only | +++<br>Myco378-3<br>only |
| Myco378-3 + GAM42a | ++*<br>Myco378-3<br>+++GAM42a | ++*<br>Myco378-3<br>+++GAM42a | +*<br>Myco378-3<br>+++GAM42a | -/+*<br>Myco378-3<br>+++GAM42a | ++<br>Myco378-3<br>only | ++<br>Myco378-3<br>only | +++<br>Myco378-3<br>only | +++<br>Myco378-3<br>only |
